# *Mycobacterium tuberculosis* FasR senses long fatty acyl-CoA through a tunnel, inducing DNA-dissociation via a transmission spine

**DOI:** 10.1101/2020.03.05.978833

**Authors:** Julia Lara, Lautaro Diacovich, Felipe Trajtenberg, Nicole Larrieux, Emilio L. Malchiodi, Marisa M. Fernandez, Gabriela Gago, Hugo Gramajo, Alejandro Buschiazzo

## Abstract

*Mycobacterium tuberculosis* is a pathogen with a unique cell envelope including very long fatty acids, implicated in bacterial resistance and host immune modulation. FasR is a two-domain transcriptional activator that belongs to the TetR family of regulators, and plays a central role in mycobacterial long-chain fatty acyl-CoA sensing and lipid biosynthesis regulation. We now disclose crystal structures of *M. tuberculosis* FasR in complex with acyl effector ligands and with DNA, uncovering its sensory and switching mechanisms. A long tunnel traverses the entire effector-binding domain, enabling long fatty acyl effectors to bind. Only when the tunnel is entirely occupied, the protein dimer adopts a rigid configuration, with its DNA-binding domains in an open state that leads to DNA dissociation. Structure-guided point-mutations further support this effector-dependent mechanism. The protein-folding hydrophobic core, connecting the two domains, is completed by the effector ligand into a continuous spine, explaining the allosteric flexible-to-ordered transition. The transmission spine is conserved in all TetR-like transcription factors, offering new opportunities for anti-tuberculosis drug discovery.

## INTRODUCTION

The composition and complexity of the mycobacterial cell envelope are two of the most distinctive features of this genus. The complex lipids present in the cell wall of *Mycobacterium tuberculosis* (*Mtb*) act as major effector molecules that interact with the host, playing key roles in pathogenicity and also providing a barrier against environmental stress, antibiotics, and the host’s immune response (1). Understanding the biogenesis of the mycobacterial cell wall will thus provide with relevant insights into the biology of this pathogen, also identifying potential targets for the development of new antimycobacterial compounds.

The outer membrane of *Mtb* comprises very long-chain fatty acids (mycolic acids), found in the inner leaflet covalently bonded to the arabinogalactan–peptidoglycan layer, and also in the outer leaflet as non-covalently associated lipids in the form of trehalose-mono- and di-mycolate (2). Mycolic acids, a hallmark of *Mycobacterium*, are synthesized by way of two fatty acid synthase systems, FAS I and FAS II. The multidomain single protein FAS I catalyses *de novo* biosynthesis of acyl-CoAs in a bimodal fashion rendering C_16-18_ and C_24-26_ derivatives (3). Long chain acyl-CoAs are used as primers by the FAS II multiprotein system, and iteratively condensed with malonyl-acyl carrier protein (malonyl-ACP) leading to very long-chain meromycolyl-ACPs (up to C_56_). The latter are eventually condensed to FAS I-synthesized C_24-26_ fatty acids (previously activated by the acyl-CoA carboxylase 4 complex (4)) to produce mycolic acids. The long-chain acyl-CoAs generated by FAS I are not only used as precursors of mycolic acids, but also for the biosynthesis of phospholipids, triacylglycerides, diverse polyketides and other complex lipids, relevant for *Mtb* pathogenicity (5,6). A complex regulatory network integrating all these pathways must exist in order to maintain lipid homeostasis. However, and despite the biological relevance of lipid-derived molecules in *Mtb*’s lifecycle, little is known about the environmental signals and the regulation cascades controlling lipid metabolism in this bacterium.

We had previously identified the transcription factor FasR, as a key activator of *fas* and *acpS* gene expression in mycobacteria (7). The *fas* and *acpS* genes form an operon, respectively coding for the FAS I synthase and the 4-phosphopantetheinyl transferase, the latter being an essential enzyme to produce functional ACP, central to all fatty-acid biosynthesis. To activate the *Mtb fas-acpS* operon, FasR binds to three inverted repeats in the promoter region, a binding that is regulated by long-chain acyl-CoAs, products of FAS I (7). More specifically, acyl-CoAs ≥ C_16_ disrupt the interaction of *Mtb* FasR with its cognate DNA. Additionally, FasR is essential for *M. smegmatis* viability (7), further highlighting its key role in mycobacterial biology.

FasR sequence reveals homology to members of the TetR family of regulators (TFRs), which are one-component sensory transduction proteins (8), typically dimeric and with each protomer displaying a 2-domain all-helical structure (9). By aligning FasR to orthologous TFRs with known 3D structures (Supplementary Fig. S1A) (10), FasR is expected to comprise a helix-turn-helix DNA-binding domain towards the N-terminus (residues 1-80), and a larger C-terminal domain corresponding to the ligand- or effector-binding domain (residues 82-228). In contrast to the DNA-binding domain, the effector-binding region reveals no sequence homology with other TFR proteins, such sequence diversity being consistent with the large variety of effector signals sensed by different TFRs (11). The prototype of the TFRs is TetR from the Tn10 transposon of *Escherichia coli*, which regulates the expression of the tetracycline efflux pump in Gram-negative bacteria (12). However, TFR proteins are widely distributed among bacteria, and control a broad range of processes, including fatty acid biosynthesis (13). Interestingly, the vast majority of TFR proteins are transcriptional repressors, with very few exceptions acting as activators (11).

We have now determined the 3D structures of FasR from *Mtb*, in three different states obtained from i- the protein crystallized alone, ii- co-crystallised in complex with the fatty acid C_20_-acyl-CoA, and iii- in complex with a double-stranded DNA oligonucleotide bearing the specific FasR-binding sequence. The comparison of these crystal structures, together with the functional characterisation of structure-guided FasR point-mutants and molecular dynamics computational simulations, uncovered the molecular mechanisms by which long- (C_16_-C_20_) and very long-chain (C_22_-C_26_) acyl-CoA molecules are sensed by FasR, as well as the means by which such signal disrupts cognate FasR-DNA binding and hence actuates *fas-acpS* transcriptional activation. World-wide efforts have disclosed hundreds of protein structures from *Mtb* corresponding to potential drug targets, a valuable input for a number of drug discovery projects (14,15). The uncovering of novel structural and mechanistic insights about a key *Mtb* metabolic regulator, contributes with solid molecular bases for target-based drug discovery, certainly one of the sensible strategies to combat tuberculosis.

## MATERIAL AND METHODS

### Bacterial strains and plasmids

*E. coli* strain DH5α cells were used for DNA cloning purposes and transformed according to standard methods. For protein expression the *E. coli* BL21 λ(DE3) strain was used instead. The *fasR* gene *(rv3208)* was PCR-amplified from *Mtb* H37Rv genomic DNA using the oligonucleotides F- Rv3208 (5’-CCCTCCATATGGAAAACCTGTACTTCCAGGGTATGAGCGATCTCGCCAAG-3’) to introduce an NdeI site at the translational start codon and encode a fused Tobacco Etch Virus protease digestion site (TEV), and R-Rv3208 (5’-GAATTCCTACGAGCGGGTAAGCG-3’) to introduce an EcoRI site at the end of the ORF. To generate a FasR recombinant protein with a hexa-histidine-TEV-tagged site at the N-terminus, the corresponding PCR product was extracted from the agarose gel, cloned into the pCR BluntII TOPO vector (Invitrogen), digested with NdeI and EcoRI, and finally subcloned into the expression vector pET28a as described by the manufacturer (Merck). The recombinant plasmid (pET28_*fasR*) was transformed into BL21 λ(DE3) cells, and transformants were selected on LB agar plates containing 50 μg/ml kanamycin. To produce the His-tagged FasR version used in all EMSA experiments (FasRwt), *fasR* was PCR-amplified from genomic DNA of *Mtb* H37Rv using the oligonucleotides F-Rv3208 (5’-CATATGAGCGATCTCGCCAAGACA-3’) to introduce a NdeI site at the translational start codon, and R-Rv3208 (5’-GAATTCCTACGAGCGGGTAAGCGG-3’) to introduce an EcoRI site at the end of the ORF. To generate a *fasR* His-tag fusion gene, the PCR product was cloned into the pCR BluntII TOPO vector, digested with NdeI and EcoRI and subcloned into NdeI/EcoRI-cleaved pET28a. The resulting plasmid (pET28_*fasR_H_*) was transformed into BL21 λ(DE3) cells. The plasmids to produce recombinant FasR_Δ33_ and FasR point mutants (FasR_LVL_, FasR_L106F_, FasR_L98A_ and FasR_F123A_) with N-terminal hexa-histidine-TEV-tagged sites, were synthesized (GenScript). NdeI/HindIII restriction sites were engineered in a pUC57-Am plasmid where genes of interest were introduced. Synthetic plasmids were digested with NdeI and HindIII, extracted from the agarose gel, and inserted into the expression vector pET28a, as described by the manufacturer (Merck). The recombinant plasmids were transformed into BL21 λ(DE3) cells and transformants selected on LB agar plates with 50 μg/ml kanamycin. DNA sequences of all genes were verified by Sanger sequencing.

### Expression and purification of proteins

Expressions of all N-terminally hexa-histidine-TEV-tagged or hexa-histidine-tagged proteins used in this work were carried out following isopropyl-β-thiogalactoside (IPTG) induction in BL21 λ(DE3) *E. coli.* Bacteria were grown at 37 C in 500□ml LB broth to 0.6-0.7 absorbance at 600□nm. IPTG was then added to 0.3-0.5□mM and the culture was grown for 12□h at 23 C. Cells were harvested by centrifugation at 2800 *g* at 4°C, resuspended in 30□ml of lysis buffer (50□mM Tris.HCl pH 8, 150□mM NaCl, 5□mM imidazole, 10 % glycerol, 10mM β-mercaptoethanol) and lysed by sonication. After centrifugation (25000 *g*, 30□min, 4 C), the supernatant was recovered and FasR, FasR_Δ33_ or FasR point mutants were separated from whole-cell lysates by Ni-NTA agarose chromatography (Qiagen, Inc). After three washing steps with lysis buffer, His_6_-tagged FasR were eluted from the resin with 250□mM imidazole in lysis buffer, dialysed overnight against FasR Buffer 1 (10□mM Tris.HCl pH 8, 300□mM NaCl). In the case of hexa-histidine-TEV-tagged proteins (FasR, FasR_Δ33_, FasR_LVL_, FasR_L1069F_, FasR_L98A_ and FasR_F123A_) after three washing steps with lysis buffer, 0.5 mg TEV protease and DTT to 1 mM final concentration were added. The mixture was incubated 2-3 hs at 23 C and 12 hs at 4 C. Proteins were eluted from the resin with 5□mM imidazole in lysis buffer and dialysed overnight against FasR Buffer 2 (10□mM Tris.HCl pH 8, 300□mM NaCl, 5% glycerol). A final size-exclusion chromatography step (S200-Superdex, GE) was performed for all proteins, pre-equilibrating the column in FasR Buffer 2 and eluting isocratically at 0.5 ml/min with high-performance liquid chromatography (Akta Purifier, GE). FasR_LVL_ was the only mutant that showed detectable levels of monomeric form eluting from SEC (all other proteins eluting as dimeric species), in which case the peak corresponding to the dimer was recovered for further functional analyses by EMSA. Protein purity was controlled by Coomassie blue staining after SDS–PAGE on a 15 % polyacrylamide gel. Protein concentrations were determined by UV spectroscopy. Purified proteins were stored at 4 C.

### Surface plasmon resonance

Surface plasmon resonance analyses were performed with a Biacore T100 instrument (GE Healthcare). Pure FasR and FasR_F123A_ were dialysed against 10 mM sodium acetate pH 4.5 and coupled to a CM5 sensor chip using the Amine Coupling Kit (GE Healthcare). Micromolar concentrations of C_20_-CoA were dialysed against 25 mM Tris.HCl pH 8, 150 mM NaCl, 0.005 % Tween 20. Serial two-fold dilutions were made in the same buffer and injected over the chip surface. Dissociation was then carried out by injecting buffer alone. Nonspecific binding was considered by injecting identical analyte concentrations over a control surface with no protein. Experiments were performed at 25 C by triplicate, producing standard deviations of less than 12 %. Data were analysed using Biacore T100 Evaluation software.

### Electrophoretic mobility shift assays (EMSAs)

His_6_-tagged FasR, and FasR point mutant proteins (FasR_LVL_, FasR_L106F_, FasR_L98A_ and FasR_F123A_) were used to assess the protein binding to P*fas*_MT_ (398 bp) promoter fragment. The promoter DNA fragment for these assays was generated by PCR amplification from *Mtb* genomic DNA with primers N2_Fas1Mt-prom (5’-CATAACGATTTGATAACAAAACTGC-3’) and C_Fas1Mt-prom (5’- CACCCGGTCGTGCTCGTGGATCGTC-3’). N2_Fas1Mt-prom primer was end-labelled with [γ-^32^P] ATP (3,000 Ci mmol^-1^) using T4 polynucleotide kinase and the PCR product obtained was purified from agarose gels. Proteins were either pre-incubated or not with acyl-CoAs and then the mixtures incubated with ^32^P-labelled probe (1000-5000 cpm) in a total volume of 25 μL binding buffer (25 mM Tris.HCl pH 8, 1 mM PMSF, 5 % (v/v) glycerol, 5 mM MgCl_2_, 150 mM NaCl and 1 μg poly-dIdC) at room temperature for 15 min. DNA-protein complexes were resolved by electrophoresis on a 6% (w/v) non-denaturing polyacrylamide gel in 1X TBE (89 mM Tris Base; 89 mM Boric Acid; 2 mM EDTA), 5 % (v/v) glycerol at 150 V, on ice. Results were visualized and recorded with a Typhoon^™^ FLA 7000 scanner (GE).

The equilibrium dissociation constant (K_D_) is a quantitative measurement to assess the affinity of biological interactions. For the FasR:DNA EMSA binding experiments described in this work we define the K_D_ as the concentration of FasR for which 50% of the DNA is in complex with the protein. The relationship between K_D_ and affinity is reciprocal, lower K_D_s correspond to higher affinities. After performing binding reactions in which the protein is titrated, the fraction of DNA bound at each concentration of protein is calculated and the data are adjusted to a binding equation using nonlinear regression (16). Densitometry of DNA bands was performed with GelPro software considering background subtraction. The fraction of bound DNA was plotted as a function of protein concentration and fit to the following binding equation (**Eq. 1**) using Prism software to perform non-linear regression:

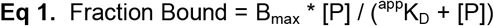

Knowing [P] = protein concentration, the apparent dissociation constant (^app^K_D_) can be quantified as well as the maximal fraction bound (B_max_) plateau.

Acyl-CoA effectors trigger FasR:DNA dissociation, and can thus be treated as non-competitive binder inhibitors. Dose-response experiments were performed to assess inhibition activity, using a modified EMSA protocol. A constant and saturating concentration of protein was equilibrated with 0.3 nM labelled DNA and increasing concentrations of acyl-CoAs in equilibration buffer. The EMSA data were collected as above and fit to a sigmoidal dose-response equation in Prism (**Eq. 2**) to determine the acyl-CoA concentration required to displace half of the bound FasR:DNA complex (IC_50_):

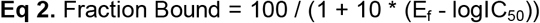

E_f_ = logarithm of acyl-CoA concentration.

EMSA experiments with FasR_wt_ and selected FasR mutants were performed by triplicate, to express ^app^K_D_ and IC_50_ as average values ± one standard error of the mean.

Apparent inhibition constants (^app^K_i_) were also calculated (17,18), to correct IC_50_ figures taking into account the K_D_ as well as the concentrations of labelled DNA ([D]) and protein ([P]) according to the Lin and Riggs conversion (**Eq. 3**):

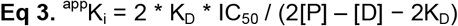

### Crystallisation and data collection

FasR_Δ33_ (5 mg/ml) crystallised at 20 C, mixing 2+2 μl of protein and reservoir solution (0.1 M MES monohydrate pH 6.0, 22 % v/v polyethylene glycol 400) using 1 ml reservoir on a hanging-drop vapour-diffusion setup. Transferred to mother liquor with 20 % (v/v) glycerol as cryoprotectant, crystals were mounted in cryo-loops (Hampton Research), and flash cooled in liquid nitrogen (19). X-ray diffraction data were collected at −163 C at the in-house Protein Crystallography Facility of the Institut Pasteur de Montevideo (Uruguay) with a MicroMax-007 HF rotating Cu anode (Rigaku) and a Mar345 image plate (marXperts).

FasR_Δ33_ (5 mg/ml) was crystallised in complex with C_20_-CoA using a 1:1 molar stoichiometry of the acyl-CoA ligand in the crystallisation drops (2 μl protein:ligand + 2 μl mother liquor 2.2 M NaCl, 0.1 M Na-acetate trihydrate pH 4.7). A hanging-drop vapour-diffusion setup was incubated at 20 C using 1 ml mother liquor as reservoir. Crystals were transferred to mother liquor with 25 % (v/v) glycerol, mounted in cryo-loops and flash cooled in liquid N_2_. X-ray diffraction data were collected at −173 C at SOLEIL synchrotron (PROXIMA 1 beamline, France) (20), using a PILATUS 6M detector (Dectris) (21).

FasR-DNA crystals were obtained by co-crystallisation. Double-stranded DNA was generated by co-incubating the two complementary oligonucleotides FwPfas25nt (5’- TACCCGTACGTAGAACTCGCCAGTA-3’) and RvPfas25nt (5’- TACTGGCGAGTTCTACGTACGGGTA-3’) under standard slow-cooling hybridisation conditions. Double-stranded DNA was mixed in a 1:1 molar stoichiometry with full-length FasR (5 mg/ml) and set for hanging-drop vapour-diffusion crystallisation by mixing 2 μl of protein:DNA with equal volume of mother liquor (28 % (w/v) PEG monomethyl ether 2000, 0.25 M ammonium citrate pH 7.0, 0.1 M imidazole) over 1ml mother liquor as reservoir solution. Crystals were transferred to mother liquor with 35 % (v/v) PEG 400, mounted in cryo-loops and flash cooled in liquid N_2_. X-ray diffraction data were collected at −173 C at Diamond Light Source synchrotron (I04-1 beamline, UK), using a PILATUS 6M detector (Dectris).

Bragg diffraction intensities were integrated with XDS (22), and scaled and reduced to amplitudes with Aimless and Ctruncate (23).

### Structure determination and refinement

The structure of FasR_Δ33_-C_20_-CoA was solved *ab initio* with Arcimboldo (24,25) which uses Phaser (26) as molecular replacement (MR) engine to place α-helices, and ShelxE (27) for density modification and chain-trace extension. The structure of FasR_Δ33_-C_14_ was solved by MR (26) using the refined FasR_Δ33_-C_20_-CoA model as search probe. Buster (28) was used to refine both FasR_Δ33_-C_20_- CoA and FasR_Δ33_-C_14_ atomic models, iterating with manual model rebuilding and validation with Coot (29). Final validation was done with MolProbity (30). OMIT maps were calculated for the FasR_Δ33_-C_20_- CoA and FasR_Δ33_-C_14_ structures according to the Polder approach (31), using final refined models from which only the acyl-CoA ligands were omitted to calculate structure factors. Final refined maps were also used to calculate real space correlation coefficients with respect to model-derived maps, on a per residue basis (32).

The FasR-DNA complex was solved by MR (26) using a portion (corresponding to the dimeric regulatory domains with no ligands nor HTH domains included) of the refined FasR_Δ33_-C_20_-CoA model as search probe. Initial refinement (33) with this partial model reduced R-factors to <50 %, and produced Fourier difference maps that clearly revealed the presence of both HTH domains and double-stranded DNA. Limited resolution and model incompleteness resulted however in mediocre 2mF_obs_-DF_calc_ maps at this point. ShelxE (27) was instrumental at improving electron density continuity, using the unrefined MR solution model and diffraction intensities as inputs, a larger than usual sphere of influence (5 Å) for density modification, the free-lunch option set at 3.85 Å to better handle data incompleteness at the higher resolution shells, and testing different solvent contents (from 0.4 to 0.6). All output maps were visualized superposed, readily allowing for manual main-chain tracing (29) of HTH and DNA portions, only including residues clearly visible in electron density at each cycle. This procedure was cycled iteratively using the progressively more complete protein models, running 5 cycles of ShelxE density modification each time. For the manual model (re)building, recent developments for real-space fitting in very low-resolution maps within Coot (34) proved essential, using Prosmart-generated external restrains, map blurring and optimized Geman-McClure parameters (alpha=0.4 proved best) for optimal movement of DNA within electron density.

The orthorhombic C222_1_ space group of the FasR-DNA crystals was confirmed by several standard procedures (23,35), notably including integration in the corresponding triclinic group and using molecular replacement as a means of deducing real symmetry. However, the DNA molecule (one double-stranded oligonucleotide per ASU) was found sitting with its long axis approximately perpendicular to one of the orthorhombic 2-fold axes, revealing both alternative 5’→3’ orientations to be present in different unit cells of the crystal. This form of static disorder is frequent when a crystal contains two or more very similar molecules (such as complementary strands in DNA with palindromic sequences), which may occupy the same lattice position with low occupancy. Refinement was performed with phenix.refine (33), using external restrains from the high resolution FasR models described above and overall strategies to deal with low resolution data. Intermediate cycles of real space refinement (36) were also instrumental. The occupancies of DNA atoms were reduced to 0.5, avoiding bumping restraints.

Difference Fourier maps very clearly revealed the positions of the two missing DNA-binding domains as well as the double-stranded DNA molecule early on during refinement. However, difference maps did not show any detectable signal at the expected positions of putative ligands bound within the effector-binding tunnel. To confirm this, the acyl moieties from the FasR_Δ33_-C_14_ model were superimposed in place within the FasR-DNA effector-binding tunnel, manually regularised to minimise clashes and added into the FasR-DNA model for further restrained refinement. Not only the crystallographic R-factors increased significantly (for both working and free sets of reflections), but difference Fourier maps revealed >5 σ negative peaks on the acyl atoms, indicating they are not actually present in the crystal.

Structural analyses were done with the CCP4 suite (37), PISA (38) and illustrations produced with Pymol (39).

### Structural bioinformatics to define the hydrophobic spine

FasR structural homologues were searched with PDBeFold (40), thus retrieving a wide range of sequence similarities. Structural alignment of such hits was performed with T-Coffee Expresso (41). The resulting 337 sequences were filtered keeping only 76 that had < 80% identity. This multiple sequence alignment (MSA) served to generate a profile hidden Markov model using HMMBuild (42). The UNIPROT database was searched with this profile HMM, CD-HIT (43) was used to filter redundant sequences, ultimately producing a list of 2591 sequences, all containing TFR effector-binding and DNA-binding domains. A MSA was calculated with T-Coffee in M-Coffee mode (44), and the resulting alignment allowed the calculation of observed frequencies for each one of the 20 amino acids for each position in the MSA. A score for each MSA position was then generated by multiplying each frequency by the corresponding hydrophobicity index (45) and summing for all 20 amino acids. The hydrophobic spine was defined as the set of positions with a weighted hydrophobicity index (which goes from −4.5 [hydrophilic] to 4.5 [hydrophobic]) equal or larger than 2. Sequence alignment figures were prepared with Espript (46).

### Molecular Dynamics simulation of FasR_Δ33_ bound to C_26_-CoA

The FasR_Δ33_-C_26_-CoA complex was built using the FasR_Δ33_-C_20_-CoA (PDB 6O6N) model as template. The bound acyl-CoA was manually extended by six carbons using Pymol (39). C_26_-CoA was optimized and 10,000 rotamers were generated with RDKit (47). Energy minimization was performed with the Rosetta suite (48) using dimer symmetry constraints, harmonic restraints were used to preserve the ligands positions as observed in the crystal structure, and 10,000 models were generated. The best complex was selected based on Rosetta energy score, with optimal stereochemical geometry and no clashes. The selected model was used as starting structure for classical molecular dynamics simulations using Gromacs 2018_cuda8.0 and GROMOS96 43a1 force field (49). An octahedron box was solvated and charge-balancing counterions were included to neutralize charges (50). Initially, the system was relaxed by energy minimization, and then equilibrated for 200 ps using a reference temperature of 27 C. Simulations were performed for 10 ns with no constraints, recording snapshots every 5 ps for analysis. Visualization of protein models and structural analyses were performed with VMD (51) and Pymol (39).

## RESULTS

### Three-dimensional structures of FasR

Recombinant FasR eluted from size exclusion chromatography suggesting a dimeric structure (~52 kDa), similar to all TFRs. Attempts to crystallise full-length FasR alone failed. We hypothesised that the 33-amino acid segment at the N-terminus is likely flexible, based on multiple sequence alignments to close orthologues (≥80% identity) from mycobacteria (Supplementary Fig. S1B). A particularly strong sequence variation of the short N-terminal extensions was revealed, also with secondary structure predicted to be absent. We thus generated a truncated form of FasR lacking the first 33 amino acids (FasR_Δ33_).

FasR_Δ33_ readily crystallised in the absence of added ligands and also in complex with acyl C_20_-CoA (arachinoyl- or arachidoyl-CoA). Both crystal forms diffracted X-rays at better than 1.7 Å resolution (Table 1), and their structures confirm the dimeric architecture of FasR_Δ33_, with each protomer organized in two all-helical domains (Fig. 1A), similar to all TFRs (11).

**Figure 1.**
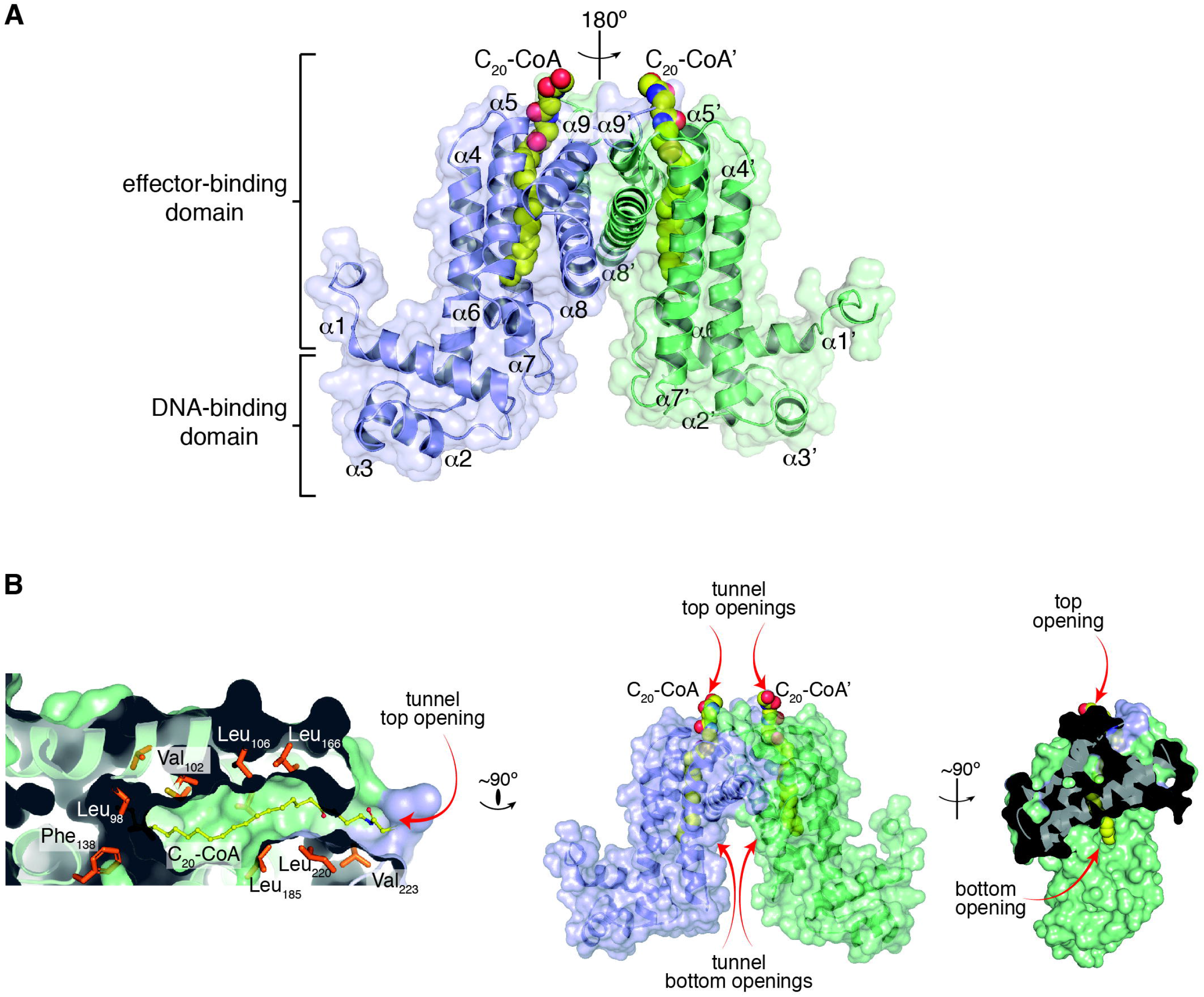
Crystal structure of FasR_Δ33_-C_20_-CoA. **A.** FasR dimer in cartoon representation, the two protomers are distinguished in light blue and pale green. The dimer is strictly symmetric, with one protomer in the asymmetric unit, the crystallographic 2-fold is depicted as a vertical axis in the plane of the figure. Secondary structure elements are labelled, a prime symbol denotes equivalent elements in the other protomer. The two domains are indicated, an N-terminal DNA-binding domain with a helixturn-helix (HTH) motif, and a C-terminal domain with the co-crystallized effector molecule C_20_-CoA bound (in spheres coloured by element). **B.** Details of the effector-binding tunnel. Three orthogonal views illustrate the molecular surface of FasR_Δ33_.The leftmost cuts along the middle of a protomer, revealing the top tunnel mouth on the right and a large segment of the tunnel itself with the bound acyl moiety (in sticks). A few amino acid sidechains that define the tunnel walls are labelled. The rightmost panel is an open-book perspective, with the footprint of protomer A visible on protomer B’s surface (in pale green), the two openings of the tunnel are visible, showing solvent exposure of both tips of the C_20_-CoA molecule.

**Table 1.**
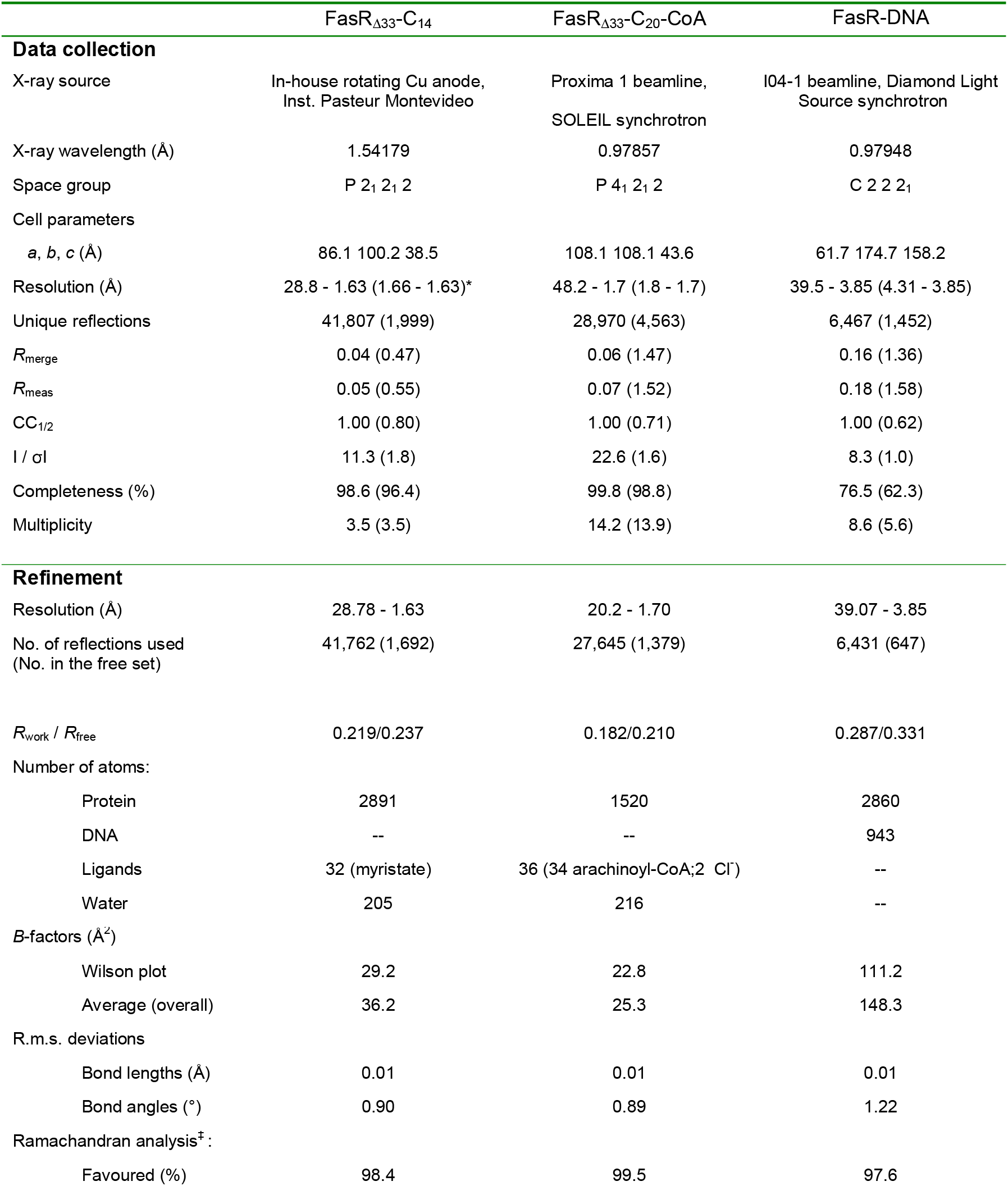
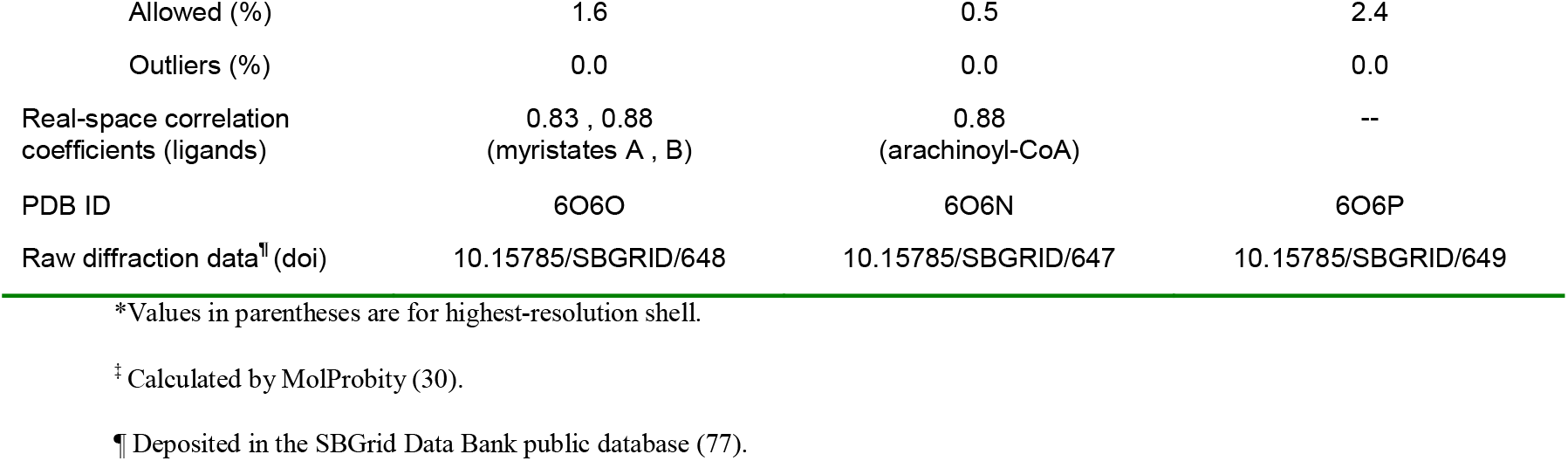
X-ray diffraction data collection and structure refinement statistics.

FasR_Δ33_-C_20_-CoA was solved first using *ab initio* methods (25), facilitated by high enough diffraction resolution and the all-helical nature of the protein. A typical TFR structural architecture was found, with a DNA-binding HTH domain comprising helices α1-α3 from the N-terminus to residue Ser_82_. And a regulatory effector-binding domain located immediately C-terminal to the HTH, from Lys_83_ to the C-terminus, including helices α4-α9. The latter are roughly organized in two bundles, the α4-α7 core runs along the long axis of the ellipsoid regulatory domain, whereas α8-α9, roughly perpendicular to the core, mediate dimerization by forming a 4-helix bundle with the other protomer’s α8’-α9’ helices (Fig. 1A). The FasR_Δ33_-C_20_-CoA dimer is strictly symmetric, with the crystallographic 2-fold axis relating one protomer to the other.

One of the most striking features of FasR_Δ33_-C_20_-CoA is a tunnel-like cavity, delimited by helices α4, α5, α7 and α8, with its two openings towards the ‘bottom’ and the ‘top’ of the regulatory domain. C_20_-CoA binds within this tunnel, in an overall parallel orientation with respect to core helices α4, α5 and α7 (Fig. 1A,B). The tunnel is ~28 Å long, with a predominance of hydrophobic residues on its wall pointing their side chains towards the lumen of the tunnel (Fig. 1B). The fatty acid is well defined all along the tunnel (Supplementary Fig. S2A). Towards the upper entrance of the cavity, most of the 4’- phosphopantetheine portion of the CoA cofactor is also observed, with the higher electron density sulphur atom being instrumental in positioning the whole C_20_-CoA moiety. Likely due to high mobility of the CoA portions exposed to the solvent, electron density is less clear towards the tip of the pantoic group, and eventually becomes indistinguishable from noise in the region corresponding to the 3’- phosphoadenosine diphosphate group, explaining why they were not included in the final refined model.

FasR_Δ33_ crystals were also grown in the absence of acyl-CoA with the goal of solving the ligand-free structure. Unexpectedly, additional electron density not corresponding to protein, was clearly visible within the FasR_Δ33_ ligand-binding tunnel (Fig. 2A, Supplementary Fig. S2B). The shape of this density is compatible with part of a polyethylene glycol molecule (PEG 400 is a component of the crystallisation mother liquor), or yet also consistent with myristic acid (C_14_). PEG is less likely, as it bears bridging oxygens along the alkyl chain, which would require satisfying H-bonding interactions in a hydrophobic environment. We hypothesize that C_14_ could have bound during protein expression in *E. coli*, and thus opted to model this fatty acid within the tunnel. Even if the chemical nature of the bound species is not certain, this piece of evidence results in two consequences: FasR_Δ33_-C_14_ is not a true apo form of the protein, but it did disclose the structure of FasR with a shorter alkyl chain bound in the effector pocket, as compared to FasR_Δ33_-C_20_-CoA.

**Figure 2.**
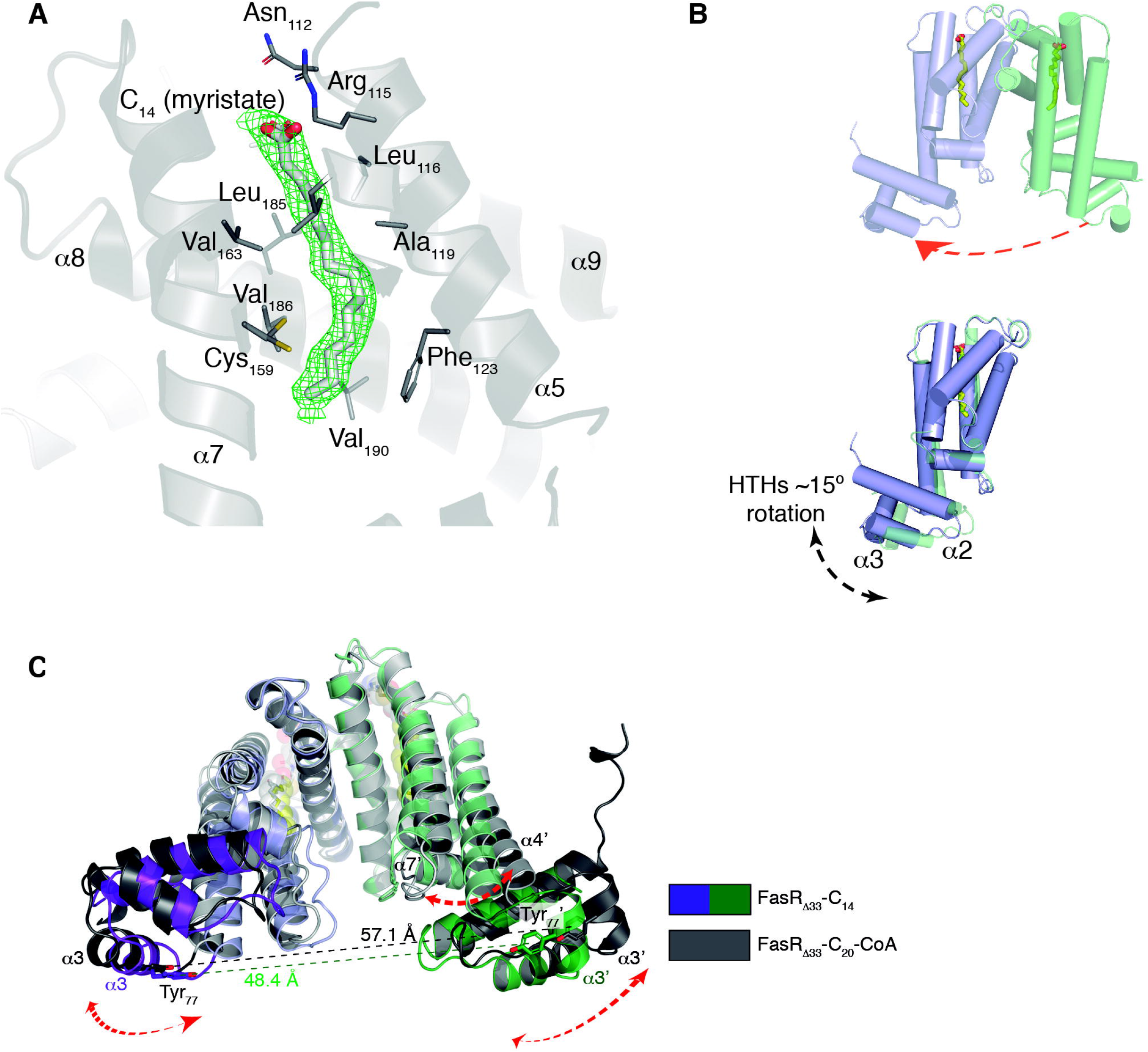
The FasR_Δ33_-C_14_ dimer is asymmetric, with larger flexibility and pendular motion of HTHs. **A.** SigmaA-weighted mF_obs_-DF_calc_ difference Fourier map contoured at 3.5 σ (green mesh) calculated with no ligand bound in the effector-binding tunnel during FasR_Δ33_-C_14_ refinement. The FasR_Δ33_-C_14_ model is shown with grey cartoons, and residues at ≤4 Å from the ligand are depicted with thin sticks. The electron density allowed to model a myristic acid, overlaid within the difference map in thick sticks coloured by atom. Note polar and charged residues on the top opening, close to the carboxyl head of the ligand. Hydrophobic residues line up the rest of the tunnel’s walls. **B.** FasR_Δ33_-C_14_ protomers A and B are coloured as in Fig. 1. The dimer is asymmetric, quantitated by superimposing protomer B onto A (red arrow in the top panel), after what the effector-binding domains fit quite well whilst the HTH domains show significant rotation between one another (bottom panel), with max root mean square deviation (rmsd) on helices α2 and α3. **C.** The FasR_Δ33_-C_20_-CoA and FasR_Δ33_-C_14_ structures are superposed, resulting in maximal fit on the central 4-helix bundle that mediates dimerization (helices α8-α9 from both protomers). The upper portion of the effector-binding domains end up well aligned, but significant shifts affect the lower part together and the HTH domains (dashed arrows). The distance between the centres of mass of Tyr_77_ on helix α3, differs by more than 10 Å between both structures. The HTH domains move consolidated with the bottom portion of juxtaposed helices (α4 and α7).

In contrast to the strictly symmetric organization of the FasR_Δ33_-C_20_-CoA dimer, FasR_Δ33_-C_14_ displays one full dimer per asymmetric unit (Fig. 2B) with each protomer deviating from strict symmetry with respect to the other. A strong 2-fold non-crystallographic operator is nonetheless relating both protomers, but applying only to the regulatory domains. The HTH domains of FasR_Δ33_-C_14_ depart from this relationship, readily observed by superimposing one protomer onto the other (Fig. 2B), which results in the regulatory domains fitting together quite well, while the DNA-binding domains appear rotated by as much as 15°. To further characterise this symmetry deviation, two identical FasR_Δ33_-C_14_ dimers were superimposed by maximizing the fit between one HTH domain from distinct protomers on each dimer (0.3Å root mean square deviation [rmsd] considering all atoms of the two superimposed HTHs). This rotation operation resulted in >5.5 Å rmsd between the other pair of HTHs. A similar exercise using the regulatory domains revealed less difference, 0.6 Å on the superimposed pair of domains *vs* 1.5 Å for the other. Departure from intradimer symmetry is thus largely due to substantial flexibility in the region that joins the regulatory and the DNA-binding domains, not the hinge loop covalently linking both domains, but rather the whole region involving the HTH as a rigid domain plus the lower part of the regulatory domain’s helices that interact with the HTH (mainly the N-terminal half of α4). Such type of “pendular” flexibility (Fig. 2B,C) is further confirmed by higher atomic displacement parameters and weak electron density in the α6- α7 loop as well as in the N-terminal half of helix α7, features that were not apparent in FasR_Δ33_- C_20_-CoA that displayed more rigidity including in the HTH domains.

The dimerization interface is an extremely well conserved structural feature among the entire TFR (11), always involving a helical bundle constituted by equivalent helices, α8 and α9 from each protomer according to the FasR helix numbering scheme. The dimers of FasR_Δ33_-C_14_ and FasR_Δ33_- C_20_-CoA were thus superimposed maximizing the fit of the dimerization helix bundles. In this way, clear differences between both structures were unveiled (Fig. 2C). In FasR_Δ33_-C_20_-CoA, the dimer is opened up, with both protomers separating away from each other, compared with FasR_Δ33_-C_14_. Using the position of conserved Tyr_77_ Cα as a reference (at the centre of helix α3, key in mediating DNA-protein binding), the two HTH domains in FasR_Δ33_-C_14_ are ~48 Å apart, *vs* ~57 Å in FasR_Δ33-_C_20_-CoA (Fig. 2C). Taking the above observations together, we hypothesize that binding of a ligand, long enough to fill up the entire hydrophobic tunnel, stabilizes the dimer in a more rigid configuration, fixing an ‘open’ HTH geometry that is not compatible with DNA-binding. In contrast, when the tunnel is occupied with a too short acyl chain, a greater mobility of the DNA-binding domains is enabled, eventually allowing the HTH helices α3 to accommodate to the required separation distance and bind DNA efficiently (ideal B-DNA has 34 Å separation between major grooves on the same side of the double-helix). This flexibility cannot be described as a simple rigid-body movement of one domain with respect to the other (a hinge-like interdomain flexibility), but rather a pendular movement implicating the DNA-binding domain and part of the effector-binding domains, as if they were acting in a consolidated way.

### Structural bases of FasR acyl-CoA-binding and effect on DNA association

To test the structure-derived hypotheses about acyl-binding and its effect on DNA-association, point-mutants were designed to block the entrance of the ligand into the hydrophobic tunnel, selecting residues that appeared to play key roles based on crystallographic evidence. Such ‘tunnel blocking’ approach was expected to abolish ligand-triggered rigidification of the protein and consequent DNA-binding hindrance. A single mutant (FasR_L106F_) and a triple mutant (FasR_LVL_) were thus constructed. The former substitutes, Leu_106_ by a phenylalanine, at a critical position that borders the entrance of the tunnel. The triple mutant adds bulky side chains not only on position 106, but also substituting Leu_185_ and Val_163_ by phenylalanines (Fig. 3A). Point-mutations did not affect the dimeric architecture of the single mutant FasR_L106F_ (Supplementary Fig. S3), and while slightly less than 50% of the triple mutant FasR_LVL_ eluted as a monomer, >50% behaved as the wild-type protein.

**Figure 3.**
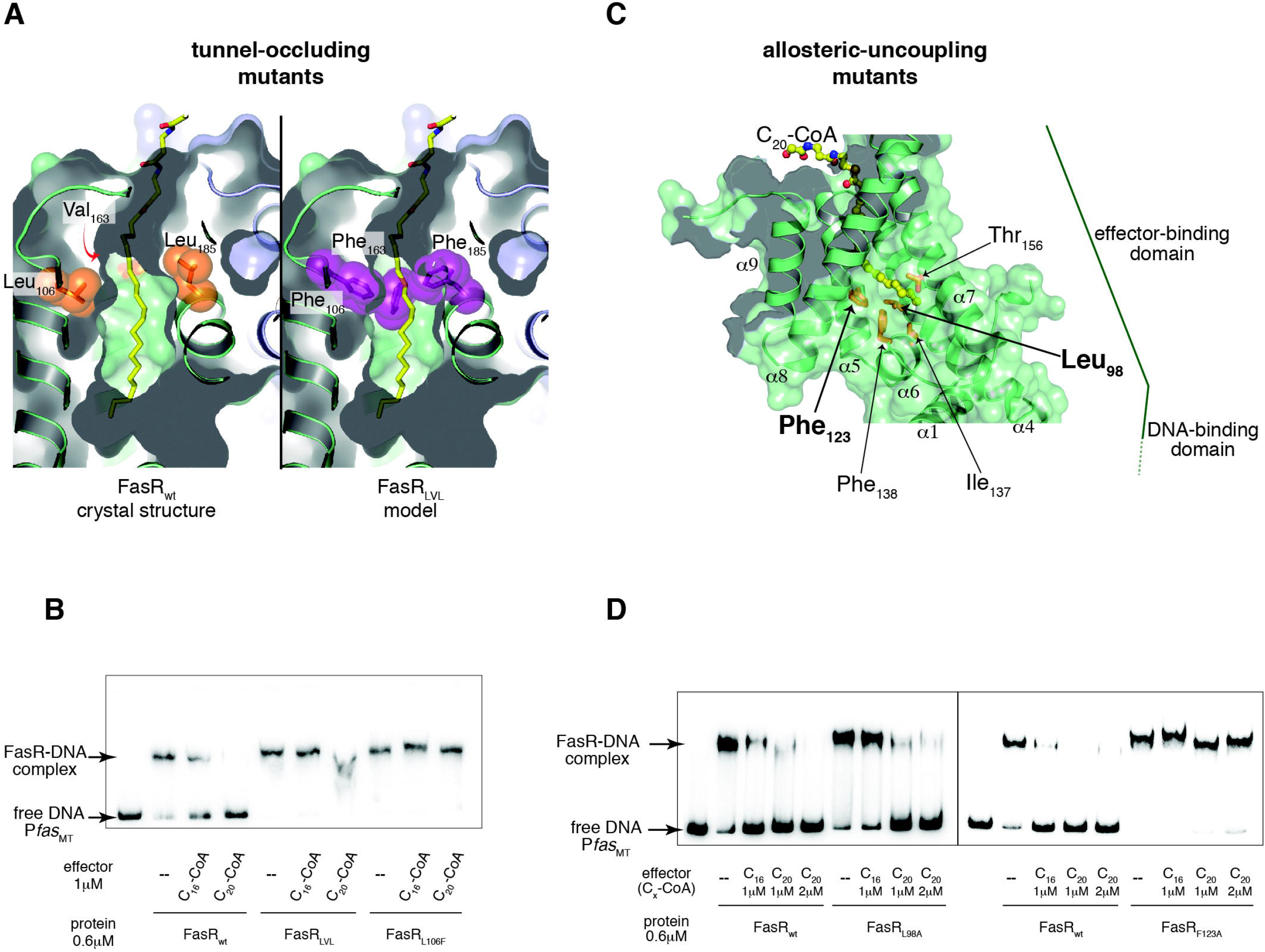
Structure guided mutations in FasR leading to permanent DNA-binding. **A**. Selected mutations to occlude the effector-binding tunnel are illustrated. The crystal structure of FasR_Δ33_-C_20_-CoA (left panel) is compared to a model of the triple mutant FasR_LVL_ (right panel). Val_163_ is actually behind the tunnel (red arrow). The substitutions by bulky phenylalanines anticipate steric hindrance of the acyl chain (shown as sticks coloured by element). **B.** Electrophoretic mobility shift assay was performed by incubating the ^32^P-labelled 398 bp P*fas* promoter region with wild-type FasR and mutant FasR proteins (FasR_LVL_ and FasR_L106F_) in the absence or in the presence of C_20_-CoA and C_16_-CoA. Protein-DNA complexes were separated by electrophoresis on a 6 % polyacrylamide gel. **C.** Selected mutations to uncouple allosteric signalling from the ligand-binding pocket to the DNA-binding domain. The bottom mouth of the effector-binding tunnel is located towards the centre of the molecule, revealing the tip of C_20_ (in ball and stick representation, coloured by atom). A few side chains are labelled and shown as orange sticks, chosen among residues bordering the tunnel opening. Two bulky residues (Leu_98_ on helix α4 and Phe_123_ on α5) that were substituted by alanines, are labelled with bold fonts. **D.** Gel shift assays of the wild-type and selected FasR mutants. The DNA probe that corresponds to the 398 bp *Pfas* promoter region (P*fas*_MT_) was ^32^P-labelled and incubated either with FasR wild-type, FasR_L98A_ or FasR_F123A_, in the absence or in the presence of the indicated concentrations of C_16_-CoA and C_20_-CoA.

Electrophoretic mobility shift assays (EMSAs) were performed by pre-incubating FasR, FasR_L106F_ and FasR_LVL_ (its dimeric form) with C_16_-CoA and C_20_-CoA, and then incubating these reactions with the ^32^P-labelled P*fas* probe (P*fas*_MT_). Strongly supporting our hypothesis, the acyl-CoA ligands triggered barely, or no detectable DNA-dissociation in the case of FasR_L106F_ and FasR_LVL_ mutants, while clearly inhibiting DNA-binding of native FasR (Fig. 3B). FasR:DNA dissociation constants (^app^K_D_) were not significantly affected by the point mutations (Supplementary Fig. S4A) and although direct monitoring of acyl-CoA binding to FasR_L106F_ and FasR_LVL_ mutants was not feasible due to technical impediments, dose-response analyses by EMSA (Supplementary Fig. S4B) further confirm that the ligands likely do not bind to the tunnel-occluding mutants. While FasR_wt_ displayed nanomolar range half maximal inhibitory concentrations (IC_50_ = 368 nM) and apparent inhibition constants (^app^K_i_ = 42 nM) of C20-CoA, the mutants exhibited significantly increased values. Namely, the FasR_L106F_ mutant only showed dissociation of the protein-DNA complex at the highest concentrations of acyl-CoA used (> 4 μM), while FasR_LVL_ remained bound to the DNA probe even at 6 μM of C20-CoA (Supplementary Fig. S4B). Ligand entrance within the tunnel is thus needed in order to trigger the protein conformational change that precludes binding of the regulator to its cognate DNA site.

### Signal transmission: allosteric mechanism connecting the ligand-binding pocket with the DNA-binding domain

The differential shift of the bottom-half of helices α4 and α7, comparing FasR_Δ33_-C_14_ and FasR_Δ33_.C_20_-CoA structures (Fig. 2C), strongly suggested that long enough alkyl chains in the effector-binding tunnel are critical in triggering the rigidification rearrangement, which results in arm-opening. To understand the molecular bases for such effect, the protein region around the distal tip of the ligand acyl chains were analysed in detail. The very last carbon atoms of C_20_-CoA interact with mostly bulky hydrophobic residues towards the end of the tunnel, *i.e.* at the opening of the tunnel that leads to the space separating both protomers in the dimer. Among these hydrophobic residues Leu_98_ (on helix α4), Phe_123_ (on α5) and Phe_138_ (on α6) might play relevant roles (Fig. 3C). The substitution of such voluminous hydrophobic residues by smaller alanine sidechains, could uncouple the HTH mobility-restraining effect from ligand-binding. Two point-mutants were constructed, FasR_L98A_ and FasR_F123A_, which maintained the dimeric architecture of the protein (Supplementary Fig. S2). Both mutants showed significant functional effects, uncoupling ligand-binding and DNA-association (Fig. 3D), with a clearer effect observed in the case of FasR_F123A_. To further dissect the underlying mechanisms, the DNA-binding affinities were first analysed, comparing wild-type *vs* FasR_L98A_ and FasR_F123A_ mutants (Supplementary Fig. S5). Apparent dissociation constants (^app^K_D_) were quantitated from electrophoretic mobility shift data, all in the nanomolar range (Table 2). Compared to FasR_wt_, FasR_L98A_ displayed slightly lower affinity for the P*fas*_MT_ probe and FasR_F123A_ higher, but neither one showed considerable effect. According to the hypothesis that these point mutations would however affect efficient transmission of the signal from effector-binding to DNA-binding domains, we next calculated IC_50_ and ^app^K_i_ values for the three proteins by evaluating dose-response effects of increasing concentrations of C_20_-CoA on protein:DNA association (Supplementary Fig. S6). Indeed, both mutants had significantly lower response to the ligand (Table 2), with a particularly pronounced effect observed for FasR_F123A_. The length of the acyl chain was also critical, with IC_50_s and ^app^K_i_s all shifted to significantly higher values when C_16_-CoA was used (Supplementary Fig. S7 and Table 2). These results strongly suggest that residues Leu_98_ and Phe_123_, and especially the latter, are key to ensure allosteric signal transmission while not influencing DNA-binding. That these mutations uncouple signal transmission but do not alter the protein’s affinity for the effector ligand was assessed for the FasR_F123A_ mutant (Supplementary Fig. S8). Surface plasmon resonance showed comparable association kinetics of FasR_F123A_ to C_20_-CoA as compared to FasR_wt_. In sum, specific residues that are not essential to bind acyl ligands nor DNA, play a key role in transmitting the signal between both domains of the protein once the effector-binding tunnel is fully occupied.

**Table 2.** Quantitative parameters of FasR association to DNA, and of its dose-dependent inhibition by effector ligands. Comparison of FasR_wt_ *vs* mutants FasR_L98A_ and FasR_F123A_ that uncouple the allosteric effect.

|  | FasR<sub>wt</sub> | FasR<sub>L98A</sub> | FasR<sub>F123A</sub> |
| --- | --- | --- | --- |
| **Binding to DNA (Pfas<sub>MT</sub>)** |  |  |  |
| <sup>app</sup>K<sub>D</sub> (nM) | 83 ± 11 | 155 ± 18 | 39 ± 6 |
| **C<sub>20</sub>-CoA dose-response** |  |  |  |
| IC<sub>50</sub> (nM) | 368 ± 105 | 634 ± 100 | 5269 ± 1067 |
| <sup>app</sup>K<sub>i</sub> (nM) | 42 | 152 | 571 |
| **C<sub>16</sub>-CoA dose-response** |  |  |  |
| IC<sub>50</sub> (nM) | 5415 ± 1047 | ND* | ND |
| <sup>app</sup>K<sub>i</sub> (nM) | 589 | ND | ND |

<sup>app</sup>K<sub>D</sub> and IC<sub>50</sub> values are reported as the average of at least three independent experiments ± one standard error of the mean.
\* ND = not detectable.

### The singular tunnel in FasR is predicted to accommodate very long fatty acyl effectors, triggering allosteric rigidification

Very long fatty acyl moieties are relevant in the biology of Mycobacteriaceae including *Mtb* (52). Intermediates in the synthesis, and constitutive moieties, of mycolic acids (with acyl components that can reach 60-90 carbons), very long fatty acids are essential components of mycobacterial cell walls. FasR activates the expression of the *fas* gene, which encodes the synthase FAS I implicated in long- and very long-chain fatty acid biosynthesis. The question is how can such long alkyl chains act as effectors of FasR? A fully extended C_20_ acyl chain measures 26.5 Å, within the FasR-C_20_-CoA tunnel the acyl shows some bending reducing that length to ~20 Å. The latter magnitude is enough for the C_20_ chain to fully occupy the tunnel, its distal tip located immediately beside the bottom opening that leads to the space between protomers in the dimer. We predict that longer acyl chains will accommodate, by extending the additional carbon atoms into the interprotomer space. To provide support for this scenario, classical molecular dynamics trajectories were calculated (Supplementary Video 1) starting from our FasR-C_20_-CoA structure where the C_20_-CoA was substituted *in silico* by the ~8 Å longer C_26_-CoA (cerotic acyl-CoA), a particularly important biosynthesis intermediate synthesized by FAS I to provide for the α-alkyl chain of mycolic acids. After initial energy minimization, 10 ns all-atom trajectories were simulated with explicit solvent, showing that the cerotic acyl chains are stable within the tunnel, their 6-carbon extensions protruding into the interprotomer space and displaying extreme flexibility (Fig. 4). The available volume in the open form of acyl-bound FasR anticipates even longer acyl chains to be readily accommodated, considering that the protein shows very stable behaviour with ~2 Å average rmsd.

**Figure 4.**
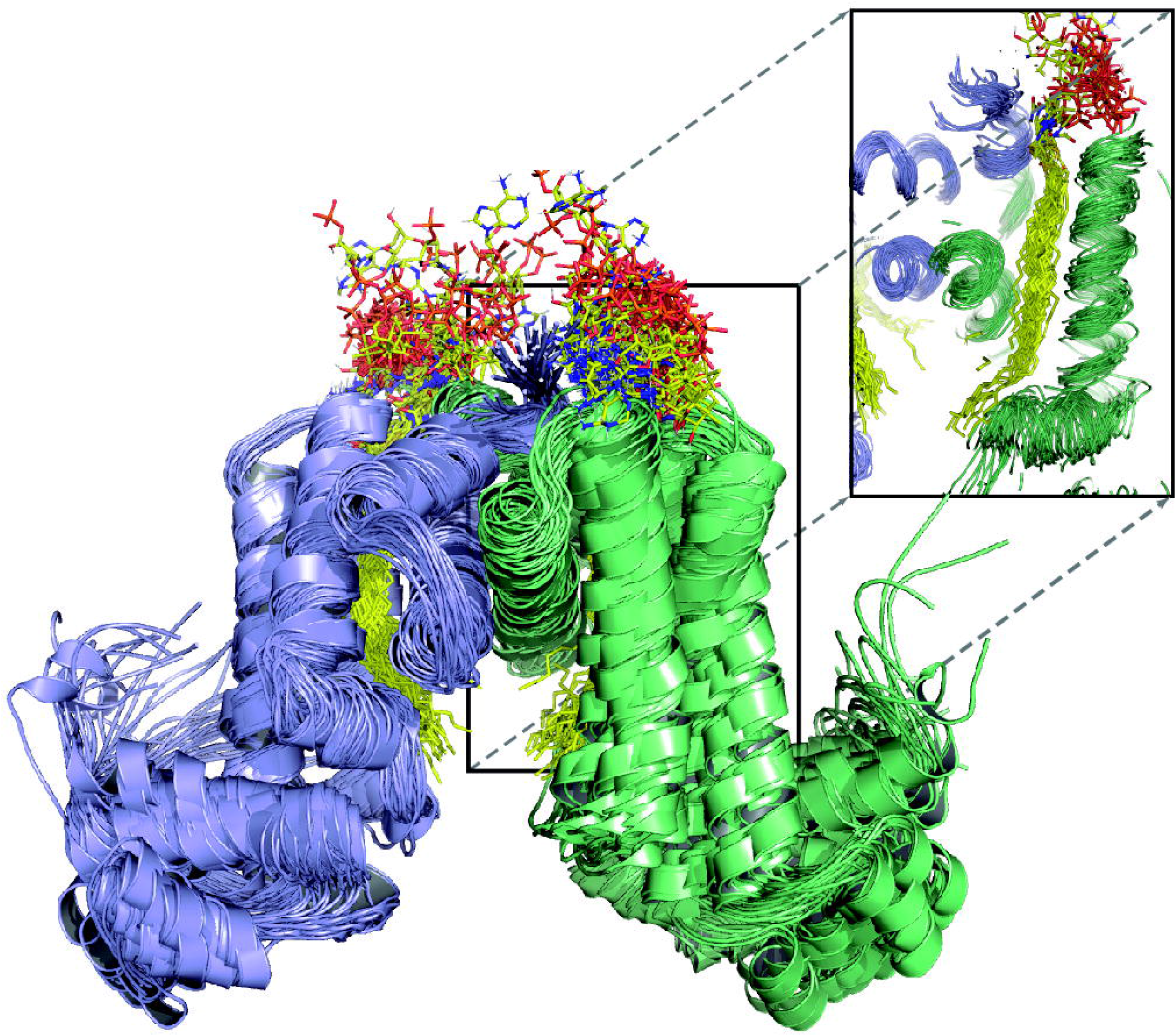
Longer than C20 acyl chains can be accommodated within the FasR_Δ33_ dimer. A representative 10ns trajectory of all-atom molecular dynamics is illustrated by showing in cartoon representation the simulated model of FasR_Δ33_ in complex with C_26_-CoA every 250 ps (see Supplementary Video 1). Shown as superposed ensemble, the secondary structure elements of both domains are preserved, and the cerotic acyl chain (in sticks) also remains stable within the tunnel. The inset shows a projection of a cut section through protomer’s B tunnel, to display the dynamic stability of the dimeric interface and the acyl chain (as opposed to the large flexibility of the nucleotide portion of coenzyme-A). Free volume is available at the inter-protomer space, predicting that even longer acyl moieties should be able to accommodate.

### The structure of effector-free FasR in complex with DNA confirms the acyl-CoA-triggered allosteric mechanism

The crystal structure of full-length FasR was eventually solved by co-crystallisation with a 25-bp double stranded oligonucleotide bearing the native FasR-binding sequence motif (Fig. 5). A number of crystals and cryo-protections methods were tested, consistently producing strongly anisotropic X-ray diffraction data, reaching 3.85 Å resolution in the best direction (Table 1). A form of crystal disorder affected the position of the DNA double helix (details in Methods), occupying equivalent positions in different unit cells while sitting alternatively in both 5’→3’ directions according to a crystallographic 2-fold.

**Figure 5.**
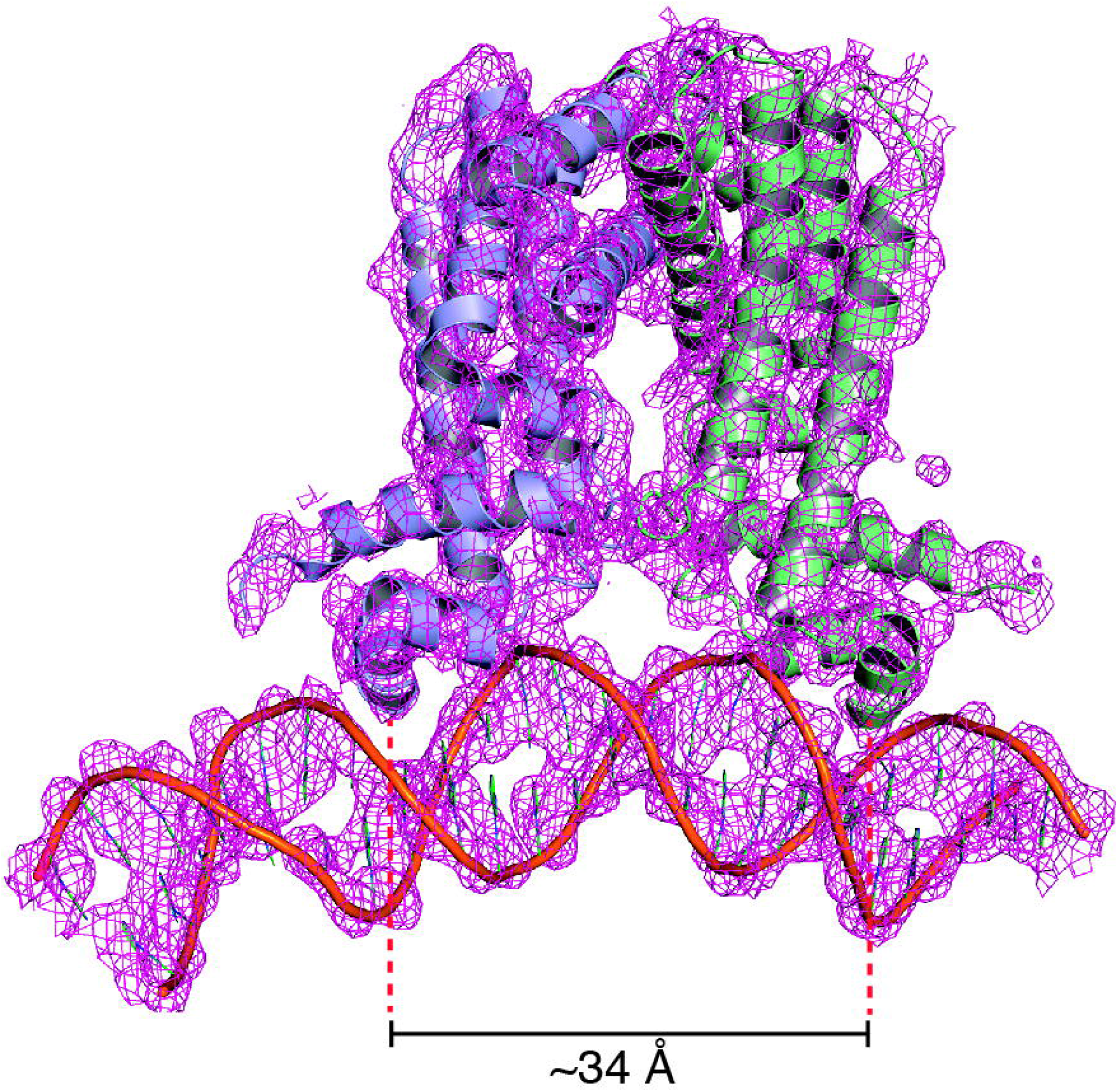
Crystal structure of full-length FasR in complex with DNA. Electron density map (sigmaA-weighted 2mF_obs_-DF_calc_ Fourier) is shown (magenta mesh), overlaid on the final refined model, showing the protein dimer (protomers in light-blue and pale-green) and the DNA double-helix (orange) in cartoon representation. The map was carved around the atomic model with a border of 3 Å to improve the clarity. Helices α3 are seen fitting within the DNA major groove as expected.

The protein that was used to grow the FasR-DNA crystals corresponds to full-length FasR. However, the first ~30-35 amino acids at the N-termini are not visible in electron density, indicating they are not bound to the DNA, likely due to high flexibility. Neighbouring FasR dimers related by crystallographic symmetry, are observed bound to the same DNA fragment on a juxtaposed consecutive manner, at roughly 90° one from the other. Such organization is associated to a pronounced bending of the DNA molecule. The position of the major kink is similar to the one identified in other TFRs, inducing a very similar DNA bending angle as in TetR (53), or yet similar in magnitude but inverted compared to the archaeal FadR (54) due to the significantly shifted positions of the HTH domains relative to the dimeric effector-binding core (Supplementary Fig. S9). The limited resolution of the FasR-DNA structure, and the presence of crystal disorder affecting the occupancy of the DNA molecules, precluded detailed analyses of protein:DNA interactions. However, the structure did provide two accurate pieces of evidence: i- the absence of ligand bound within the tunnel of the effector-binding domain; and, ii- a major conformational rearrangement bringing the HTH DNA-binding domains closer together in the dimer (Supplementary Video 2) such that helices α3 now fit within two successive major grooves on the DNA molecule.

This conformational change hampers effector occupation within the tunnel, the latter seems to be constricted by shifted residues (e.g. Phe_123_) which move their side chains towards the tunnel’s lumen. The FasR_Δ33_-C_14_ structure described above proved that acyl-containing compounds from *E. coli* are invariably associated within the effector-binding cavity, even if specific acyl/acyl-CoA molecules are not added during protein purification and crystallisation. That the FasR tunnel in the FasR-DNA complex is free of bound ligands is supported by unequivocal evidence from difference Fourier maps at both early and late stages of refinement (detailed in Methods). The association to DNA thus correlates to expelling ligands from the effector-binding tunnel of FasR, at least those that attach more loosely within the cavity.

## DISCUSSION

FasR is a TFR (**T**etR **f**amily of **r**egulators) member that senses long and very long acyl-CoA moieties, subsequently turning off FAS I-mediated fatty acid biosynthesis. *Mtb* is able to synthesize very long acyl-CoA intermediates (e.g. in the way to synthesizing mycolic acids), spanning molecular lengths of 40-50 Å and more. It is known that TFRs possess pockets, sometimes even deeper tunnel-like cavities, which bind to, and enclose the sensed effector ligand. How can FasR deal with the very long effector molecules it senses? It must be stressed that the entire effector-binding domain of FasR measures ~40 Å along its longest axis (substantially less considering inner cavities); hence, the fatty acyl effectors can often be longer than the protein’s own physical boundaries. We now answer to this question by revealing a unique hydrophobic tunnel that cuts across the entire effector-binding domain of FasR (Fig. 1), a tunnel that is conspicuously opened on both ends. This singular solution has evolved to lodge the kind of acylated chains of 20 and more carbons that inhibit FasR binding to its cognate DNA (7).

In addition to the tunnel itself, and in continuity with the bottom opening of it, FasR possesses a cavity delimited by the two protomers. The volume of such cavity (55) in FasR is unusually large (~1800 Å^3^) compared to many other TFRs: ~600 Å^3^ (*S. enterica* RamR PDB 3VVY; and *M. tuberculosis* EthR PDB 5NIO), ~530 Å^3^ (*E. coli* RutR PDB 4XK4), ~340 Å^3^ (*E. coli* TetR PDB 2XPW), ~135 Å^3^ (*S. acidocaldarius* FadR PDB 6EL2), or yet 40 Å^3^ (*P. aeruginosa* DesT PDB 3LSJ). This feature is likely relevant, as it anticipates FasR’s ability to accommodate very long fatty acyl chains, such as C_26_-CoA (synthesized by the *Mtb* FAS I system (56)), while maintaining a stable, open configuration (Fig. 4, Supplementary Video 1). Other fatty acid-sensory TFRs fairly similar to FasR (54,57-59), either do not create true continuous tunnels (58), or engage a different set of residues running in a perpendicular direction as compared to FasR’s cavity (54,57,59). EthR is yet another TFR from *Mtb* (60), intensively investigated as a target to develop anti-tuberculosis medicines. However, EthR is substantially different from FasR (22 % sequence identity; ~4.5 Å rmsd after superposition of the effector-binding domains), with a shorter N-terminal extension before the first α-helix, and a tunnel displaying wider regions or bulges (Supplementary Fig. S10). Such bulges seem to correlate with EthR’s capacity to bind compounds that include 1 to 3 aromatic or aliphatic rings (61). At difference with FasR, the physiologic molecules sensed by EthR remain unidentified, despite the >70 EthR crystal structures available, most in complex with surrogate ligands. Among these, only one corresponds to a linear chain (hexadecyl octanoate, in PDBs 1U9N and 1U9O), in this way the most similar to FasR effectors. This ligand is positioned in EthR in a similar configuration as the acyl ligands in FasR, but leaving part of EthR’s available tunnel volume unoccupied (Supplementary Fig. S10C), suggesting that physiologic effectors are likely to be of larger size, branched and/or containing bulkier cyclic groups.

FasR is thus equipped to binding very long-chain acyl effectors, but how is such binding coupled to inhibiting DNA-association? The type of rearrangements that we have found (Supplementary Video 2) are consistent with the ones observed in a number of other TFRs, in principle conforming to the mechanistic hypothesis that the binding of effector ligand (the signal) induces an HTH-open conformation, eventually inhibiting TFR association to DNA (the output response). Association of TFRs to DNA indeed requires the HTH domains to close in, in order for the α3 helices of the two protomers to fit into two successive major grooves (~34 Å apart) on the same side of the cognate DNA (62). Such a mechanism has been put forward to explain the workings of *E. coli* TetR when binding tetracycline (63), DesT from *Pseudomonas aeruginosa* sensing saturated *vs* unsaturated acyl-CoAs (58), RutR recognizing uracil in *E. coli* (64) or yet the multi-drug binding protein QacR from *Staphylococcus aureus* (65), among many others.

Binding the effector stabilizes an open configuration of FasR, which does not necessarily imply that the effector mechanically triggers a closed to open transition. To ascertain the latter mechanism, the ligand-free structure should exhibit a closed, DNA-binding competent configuration. A ligand-free form of FasR could not be crystallized, but turning our attention to the many other available TFR structures, those that exhibit no ligand almost exclusively correspond to the open form (9,11), contradicting the predicted outcome. The very few apo structures with a closed configuration have unexplained density within the binding pocket (pdb 2FX0) and/or reveal crystal packings that fortuitously fix HTH domains strongly in place (e.g. the hexagonal form of 2FX0 and the centred monoclinic 1T33). To the best of our knowledge, the closed configuration has only been captured reliably in crystal structures of TFRs in complex with DNA. Additional evidence further challenge a simple open/closure mechanism: *i-* no simple positional shifts of individual residues can explain the mechanical bases of the alleged pendular movement; *ii-* a number of TFR mutants have been identified that either uncouple effector-binding from transcriptional induction (66), or invert the effector’s action by triggering a tighter binding to DNA (67,68), in both cases often implicating positions that are not directly involved in effector-binding.

Finite deformations physics theory seems attractive to highlight allosteric regulation pathways by measuring mechanical strain rather than pairwise atomic position deviations (69). Unexpectedly, one of the segments subjected to highest mechanical strain in all TFRs analysed, corresponds to the loop that connects helices α6 and α7 (Supplementary Fig. S11). A triangle defined by α5, α6 and α7, a conserved feature in all TFRs (9), harbours the ligand-binding core cavity (which can in some cases expand into tunnel-like architectures with top, bottom and/or lateral openings). Helix α6, associating to helix α8, is attached to the fixed core, upper-half of the effector-binding domain; but simultaneously, α6’s C-terminal tip and the α6-α7 junction also associate to the moving HTH. In sum, α6-α7 loop is bound to fixed and to moving parts, eventually leading to local deformation. Residues that could explain this strain, in contact with helices α6, the α6-α7 loop and the HTH domain, led us to identifying an array of hydrophobic residues, highly conserved among TFRs (Supplementary Fig. S1 and Supplementary Data 1) and configured in three-dimensions as a continuous spine connecting the two domains of FasR. This spine belongs to, and connects the hydrophobic protein-folding cores of both domains, being interrupted by the ligand-binding cavity in all TFRs analysed (Fig. 6 and Supplementary Fig. S12). Only in the ligand-bound condition this hydrophobic spine is completed, by the ligand molecule itself at the effector-binding domain, stabilizing a rigid and open conformation. Such a mechanism predicts a disordered (flexible) to ordered transition of the TFR protein, which is consistent with the evidence we provide for FasR as well as with available evidence from other TFRs (9,11,66-68,70). In particular, fluorescent probes such as 1-anilino-naphthalene-8-sulfonate that bind to partially folded proteins in “molten globule” states (71), have been shown to bind promiscuously to apo TFRs (9). Also, effector-triggered appearance of folding cooperativity between both domains, as well as proteolysis-resistance, have been reported in wild-type but not in allosteric-uncoupled mutants of TetR (70,72). Taken together, the extensive body of evidence lends strong support to the transmission spine mechanism as the most consistent interpretation of the effector-mediated allosteric control of TFRs’ DNA-binding function.

**Figure 6.**
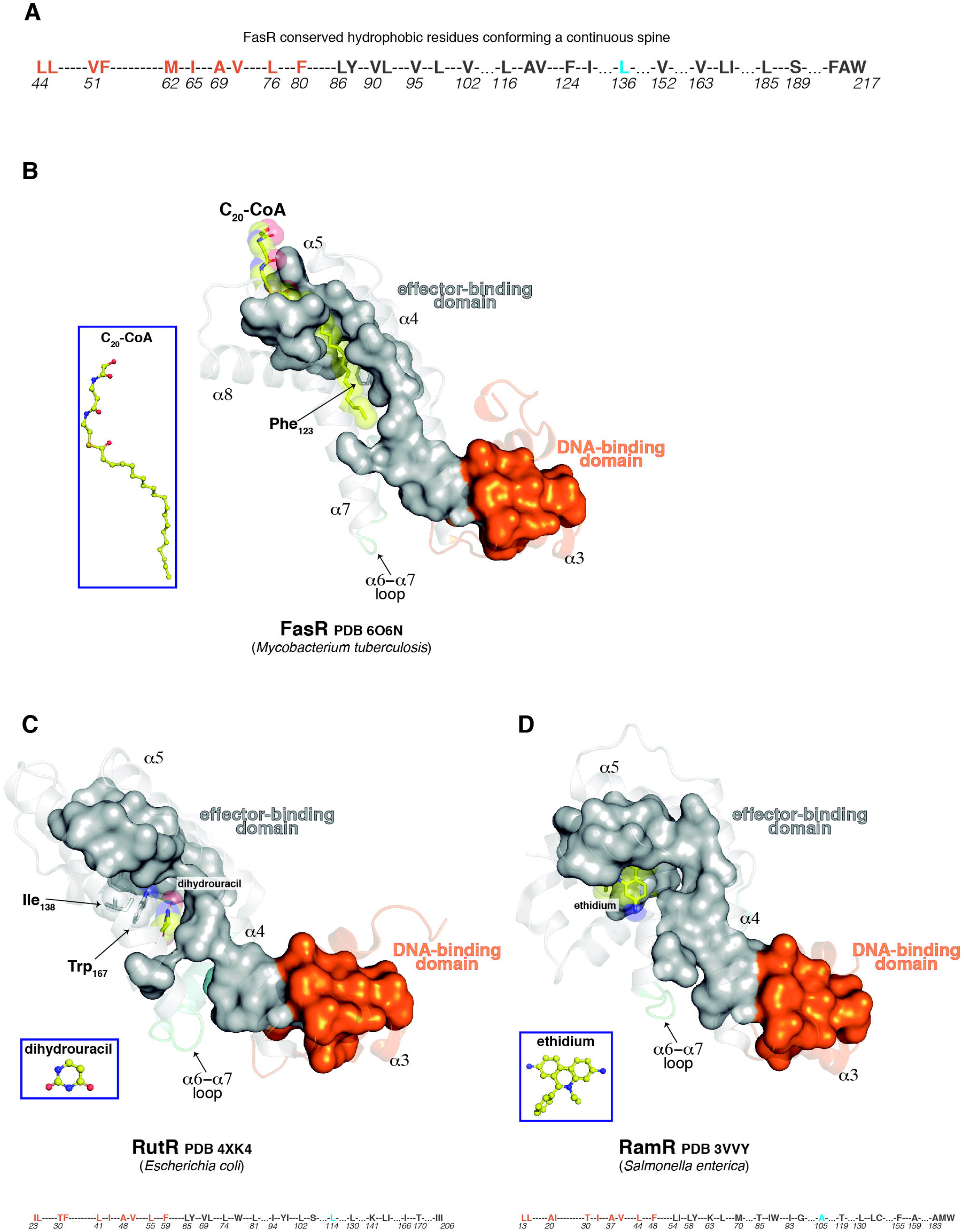
A TFR-conserved hydrophobic spine connects effector-sensing to DNA-binding output regulation. **A.** Hydrophobic spine amino acids in FasR. As detailed in Material and Methods these residues were selected after multiple sequence alignment of >2500 TFRs comprising a large sequence diversity. Thirty-four amino acid positions surpassed a global hydrophobicity cutoff, among which 32 configure a continuous spine in 3D. The colour code depicts residues of the DNA-binding domain in orange, the effector-binding domain in grey and the α6-α7 loop, subjected to strain, in cyan. **B-D.** Distinct TFRs (FasR, RutR, RamR) illustrate the conservation of the hydrophobic spine (the molecular surface of the residues is shown), connecting effector- to DNA-binding domains. Note how the binding of the effector completes the spine in every case. The colour code is identical to panel A. The effector molecules in atom-coloured sticks are labelled, with transparent spheres overlaid (insets show their markedly disparate structures). In FasR and RutR, additional residues are indicated (shown with sticks overlaid by transparent spheres), which are not included in the globally conserved spine, but that contribute to it in individual TFRs, adapting to singularities of the specific effectors. The specific sequence of the spine for each protein is shown at the bottom of panels C and D, numbered according to each protein’s sequence (colours respect the domains’ scheme). See Supplementary Fig. S12 for additional examples of TFR spines.

When the ligand leaves the site, or if it is too short to fully occupy it, the hydrophobic spine is broken, protein folding is sub-optimal, anticipating a multitude of conformations (with HTH wiggling, illustrated schematically in Fig. 7), including those that are competent for DNA-binding (HTH-closed). The disorder-to-order transition is not fully triggered, resulting in asymmetry and higher flexibility as observed in the FasR_Δ33_-C_14_ complex. This hypothesis also explains why mutating bulky residues that contribute to building and stabilising the spine (*e.g.* Leu_98_ and Phe_123_ in FasR), can uncouple the allosteric effect: the ligand is then insufficient to achieve a complete, compact fold in the mutated TFR (Supplementary Fig. S13). Residues that are not directly involved in effector-binding, but that contribute to building and stabilising the spine, will also be able to exert notable effects on allosteric coupling, upholding reported results (66-68).

**Figure 7.**
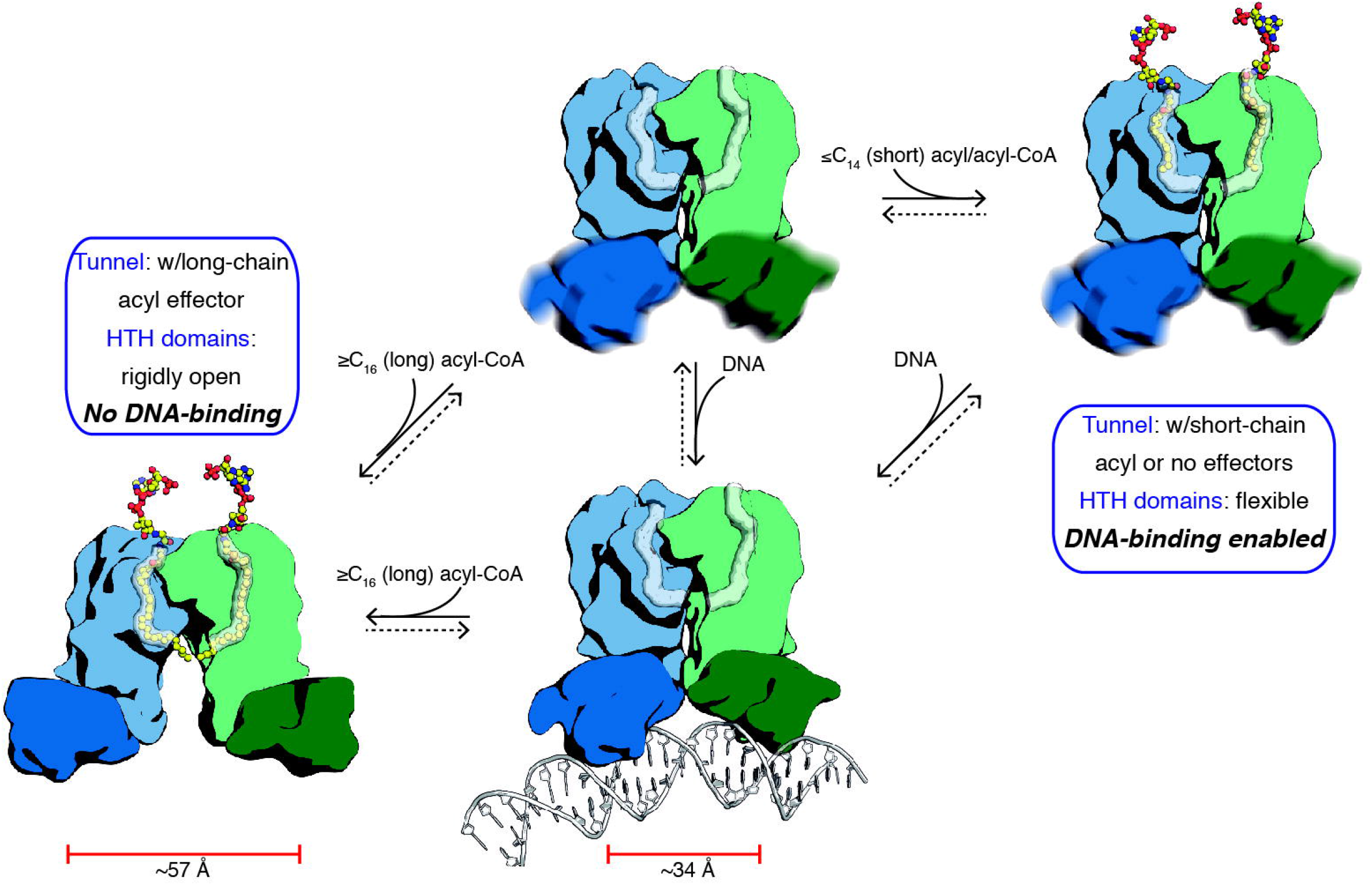
Schematic of the mechanism governing FasR-mediated long acyl-CoA sensing and response. The model combines information from the three crystal structures presented in this report (FasR_Δ33_-C_14_, FasR_Δ33_-C_20_-CoA and FasR-DNA complexes). A free structure of FasR with no bound effectors nor DNA has not been determined experimentally, and is not currently known whether it builds up to detectable concentrations within the living cell. The dotted arrows reflect plausible equilibria. See Supplementary Video 3 to better grasp the anticipated dynamics.

A second conformation, the one bound to DNA, is however compactly folded. In this case it is the polynucleotide that pulls on the flexible HTH domains of the dimer, bringing them closer together. Correlated to this HTH movement and again transmitted through the hydrophobic spine, several of the bulky residues that line up the effector-binding cavity walls, occlude the cavity (such as Phe_123_ in FasR-DNA), completing the spine into a “folded protein-like” core. This occlusion of the ligand-binding pocket in other TFRs when bound to DNA has been described in crystal structures of TetR (65), DesT (58) and FadR (59).

This scenario opens up exciting avenues to be explored. How reversible is the binding of effector compounds and what triggers their dislodging from the cavity? Following on our proposal, DNA-binding should likely expel the effector and vice-versa, depending on relative DNA-vs-effector concentrations and affinities. Of note, the assistance of other proteins, in burying the long alkyl moieties when expelled from FasR and other lipid-sensing TFRs, should not be ruled out, as they might be extremely important as part of the thermodynamic equations. The hydrophobic transmission spine hypothesis (Fig. 7) may prove instrumental to design better drugs. Eukaryotic protein kinases (ePKs) are not homologous to TFRs, but do exhibit an analogous hydrophobic spine that regulates the kinase activation switch (73), a mechanism that has been successfully exploited to develop ePK inhibitor-based drugs against human cancer and inflammatory diseases (74,75). A closer example is provided by EthR, a TetR-like repressor that controls the expression of EthA (the enzyme that bioactivates ethionamide, a second-line antituberculosis drug that inhibits mycolic acid biosynthesis). EthR inhibitors have been conceived after comparing EthR crystal structures obtained in complex with larger and smaller ligands (76). In that work, compounds bearing thienyl and piperidinyl pharmacophores were selected as the most potent EthR inhibitors among screening hits. Only the piperidinyl-, and not the thienyl-, interacting pocket engages EthR amino acids that compose the hydrophobic spine (PDBs 3G1O, 3G1M), the transmission mechanism thus explains why the piperidinyl-binding pocket turned out to be the crucial region determining inhibitory activities (76). Structure-guided drug discovery strategies that exploit the allosteric hydrophobic-spine transmission mechanism of TFRs might thus prove successful in developing novel medicines against tuberculosis, including multi- and extensively drug-resistant strains.

## AVAILABILITY

Worldwide Protein Data Bank provides open access of structural biology data to the public domain (http://www.wwpdb.org/). SBGrid Data Bank is an open source research data management system enabling Structural Biologists to preserve x-ray diffraction data and to make it accessible to the broad research community (https://data.sbgrid.org/).

## ACCESSION NUMBERS

Macromolecular 3D structural data (model coordinates and crystallographic structure factors) presented in this study have been deposited in the wwPDB with accession codes 6O6O, 6O6N and 6O6P. Raw X-ray diffraction data for each one of those structures have been deposited in SBGrid Data Bank with Digital Object Identifiers 10.15785/SBGRID/648, 10.15785/SBGRID/647 and 10.15785/SBGRID/649, respectively.

## SUPPLEMENTARY DATA

Supplementary Data are available at NAR online.

## ACKNOWLEDGEMENT

We thank Stanislas Leibler, Michael Mitchell and Pablo Sartori for sharing initial strain analysis scripts; Matias Machado for assistance in molecular dynamics; Frank Lehmann for initial cloning efforts; Sebastian Klinke (Fundación Leloir) and the staff at Proxima 1 beamline (Soleil synchrotron) and at I04-1 beamline (Diamond synchrotron) for assistance with data collection. We acknowledge computational and storage services (TARS cluster) provided by the Institut Pasteur IT Dept (Paris). We thank the CCP4/CeBEM Macromolecular Crystallography School (USP@São Carlos, 2018), especially Isabel Usón and Paul Emsley for helping us respectively with ShelxE and Coot, in dealing with low-resolution density modification and model building.

