## Supplementary Figures for "*Mycobacterium tuberculosis* FasR senses long fatty acyl-CoA through a tunnel, inducing DNA-dissociation via a transmission spine"

**Supplementary Material (Lara J *et al.*)**

List of included Supplementary Figures, Videos, Data and References

- Supplementary Figure S1
- Supplementary Figure S2
- Supplementary Figure S3
- Supplementary Figure S4
- Supplementary Figure S5
- Supplementary Figure S6
- Supplementary Figure S7
- Supplementary Figure S8
- Supplementary Figure S9
- Supplementary Figure S10
- Supplementary Figure S11
- Supplementary Figure S12
- Supplementary Figure S13
- Supplementary Video 1 (submitted as a separate MPG file)
- Supplementary Video 2 (submitted as a separate MPG file)
- Supplementary Video 3 (submitted as separate MPG file)
- Supplementary Data 1 (submitted as a separate ASCII Text file, in .mfa Multiple Fasta format for sequence alignment encoding; it can be viewed and manipulated directly by JalView or similar software)
- Supplementary References

A

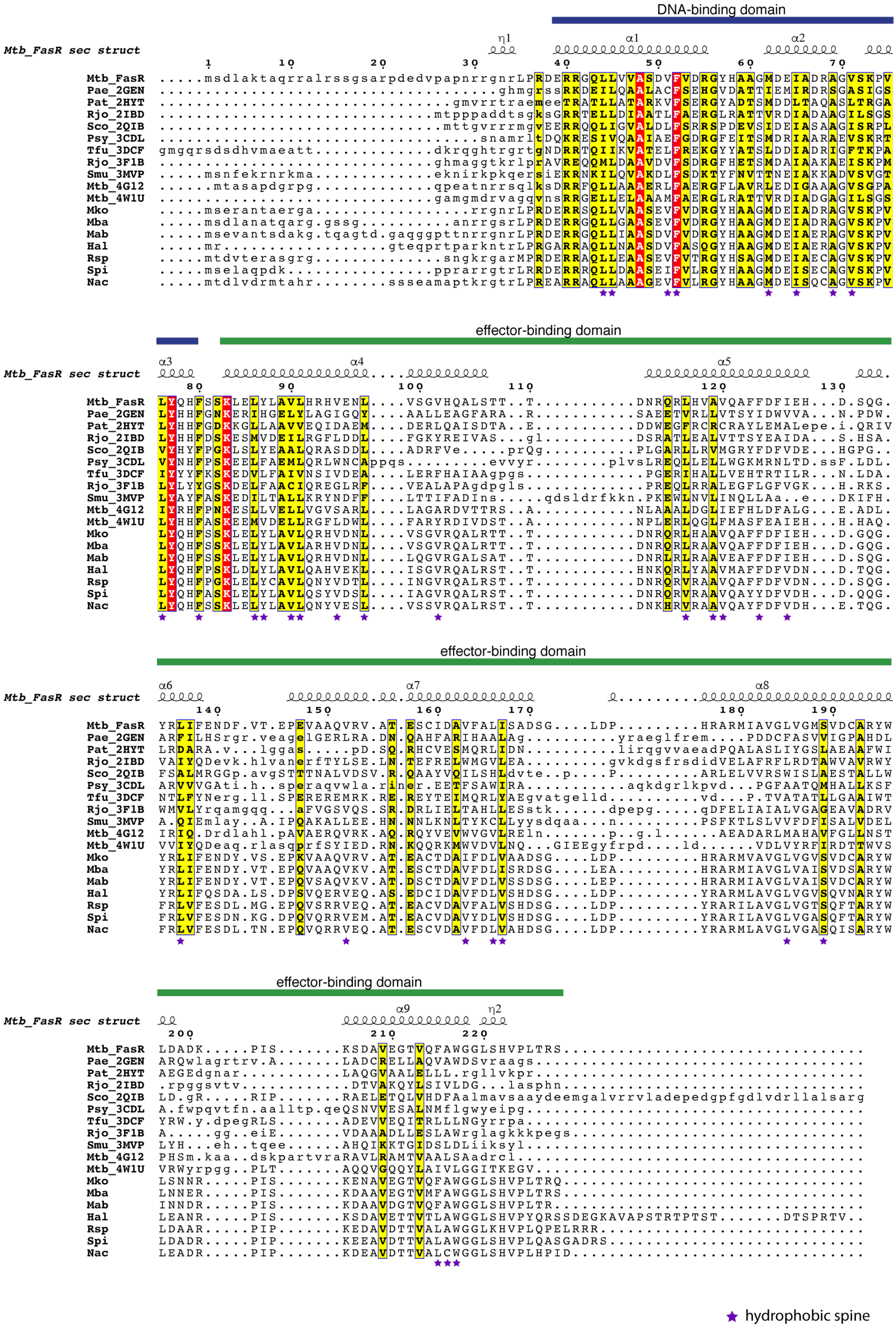

DNA-binding domain
effector-binding domain
effector-binding domain
effector-binding domain
Mtb_FasR sec struct
Mtb_FasR
Pae_2GEN
Pat_2HYT
Rjo_2IBD
Sco_2QIB
Psy_3CDL
Tfu_3DCF
Rjo_3F1B
Smu_3MVP
Mtb_4G12
Mtb_4W1U
Mko
Mba
Mab
Hal
Rsp
Spi
Nac
hydrophobic spine

**B**

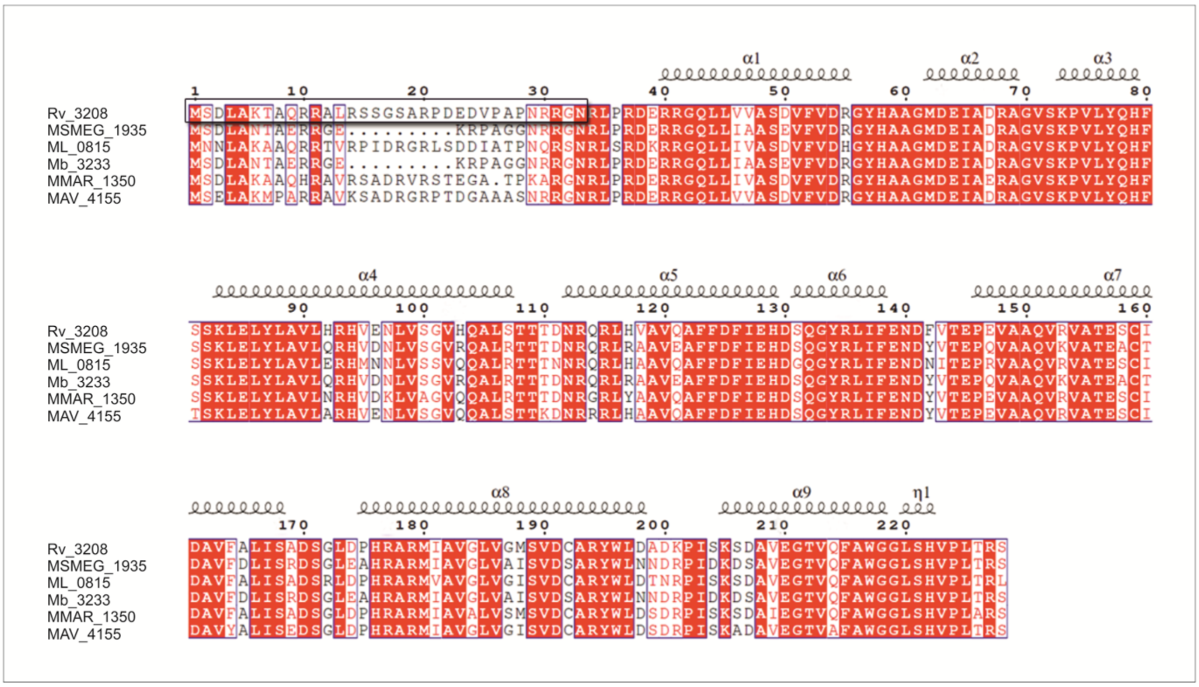

**Supplementary Figure S1**. **A.** Multiple sequence alignment of FasR (Mtb_FasR) with similar orthologues within the Protein Data Bank (orthologue sequences were chosen corresponding to the ten most similar ones on the basis of 3D structure, once FasR structure was solved, see further below in this report). The first three letters correspond to the species names, followed by the PDB code of the 3D structure. The secondary structure elements of Mtb_FasR are depicted towards the top of the alignment blocks. Purple stars indicate residues belonging to the hydrophobic transmission spine (see further below in this report). Within sequences corresponding to PDB structures, capital *vs* small letters distinguish regions that respectively can and cannot be structurally aligned with FasR. Similar residues (>0.7 global similarity score according to a Risler substitution matrix) are written with black bold characters and boxed in yellow. Invariant residues are written with white bold characters and boxed in red. The two-domain architecture of the TFR family is highlighted with blue/green coloured boxes. Seven additional sequences are included beneath the ones from the PDB (denominated with three letters corresponding to the species names) chosen among the ones with most similar DNA-binding domain sequences. Pae: *Pseudomonas aeruginosa*; Pat: *Pectobacterium atrosepticum*; Rjo: *Rhodococcus jostii*; Sco: *Streptomyces coelicolor*; Psy: *Pseudomonas syringae*; Tfu: *Thermobifida fusca*; Smu: *Streptococcus mutans*; Mtb: *Mycobacterium tuberculosis*; Mko: *Mycolicibacillus koreensis*; Mba: *Mycobacteriaceae bacterium*; Mab: *Mycobacteroides abscessus*; Hal: *Hoyosella altamirensis*; Rsp: *Rhodococcus* sp.; Spi: *Skermania piniformis*; Nac: *Nocardia acidivorans*. **B.** Sequence alignment of selected *Mycobacterium* proteins orthologous to FasR. Multiple sequence alignment of FasR Rv3208 *(M. tuberculosis)* with its orthologues MSMEG_1935 *(M. smegmatis),* ML_0815 *(M. leprae),* Mb_3233 *(M. bovis),* MMAR_1350 *(M. marinum)* and MAV_4155 *(M. avium)*. Residues that are invariant across proteins are shown in white on red background and those strongly conserved are shown in red. The remaining residues are in black. Secondary structure elements present in FasR (according to its 3D structure presented further below in this report) are shown above the alignment. The black box towards the N-terminus indicates the 33 amino acids not included in the $FasR_{\Delta 33}$ construct used for crystallographic studies. The alignment was calculated integrating structural alignment data with MAFFT (1), and the figure prepared with ESPRIPT (2)

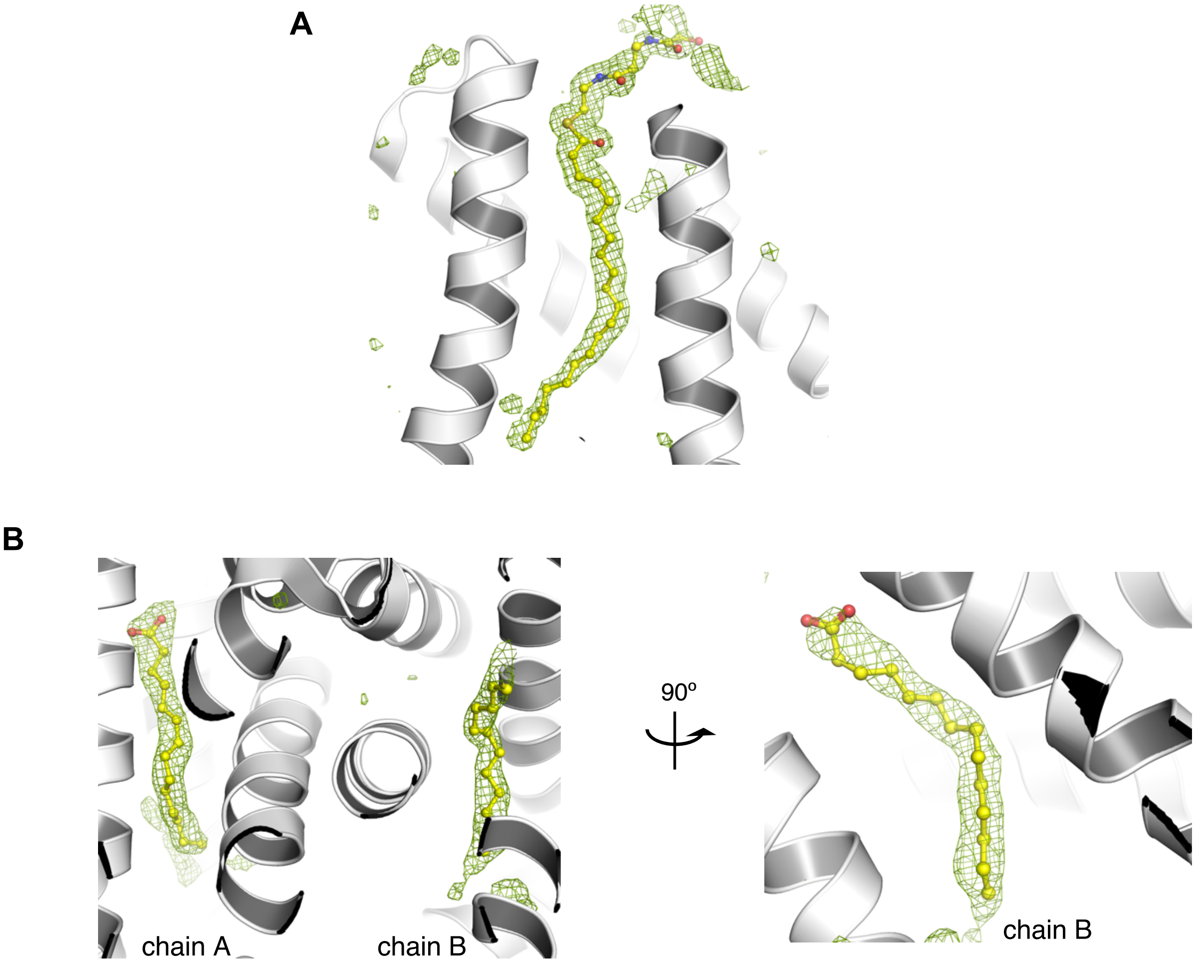


**Supplementary Figure S2. OMIT maps for ligands.** OMIT Fourier maps were calculated with [$mF_{obs}$ – $DF_{calc}$] coefficients, using calculated structure factors from the final refined models of each crystal structure, from which the corresponding acyl moieties had been deliberately omitted. **A.** OMIT map of FasR$_{\Delta33}$-$C_{20}$-CoA from which the arachinoyl-CoA ($C_{20}$-CoA) moiety was omitted. The map is contoured at 3.5σ and depicted as a green mesh. The final model of the $C_{20}$-CoA residue is superimposed in sticks-and-balls, coloured by atom. **B.** OMIT map of the FasR$_{\Delta33}$-$C_{14}$ dimer, from which the two myristate ($C_{14}$) moieties were omitted. The map is contoured at 3.5σ and depicted as a green mesh. The two monomers are labelled with their chain names. The left panel's perspective shows chain A's myristate electron density more clearly. To the right, the model was rotated 90º to render a clearer picture of the second myristate electron density. The final model of the $C_{14}$ residues are superimposed in sticks-and-balls, coloured by atom.

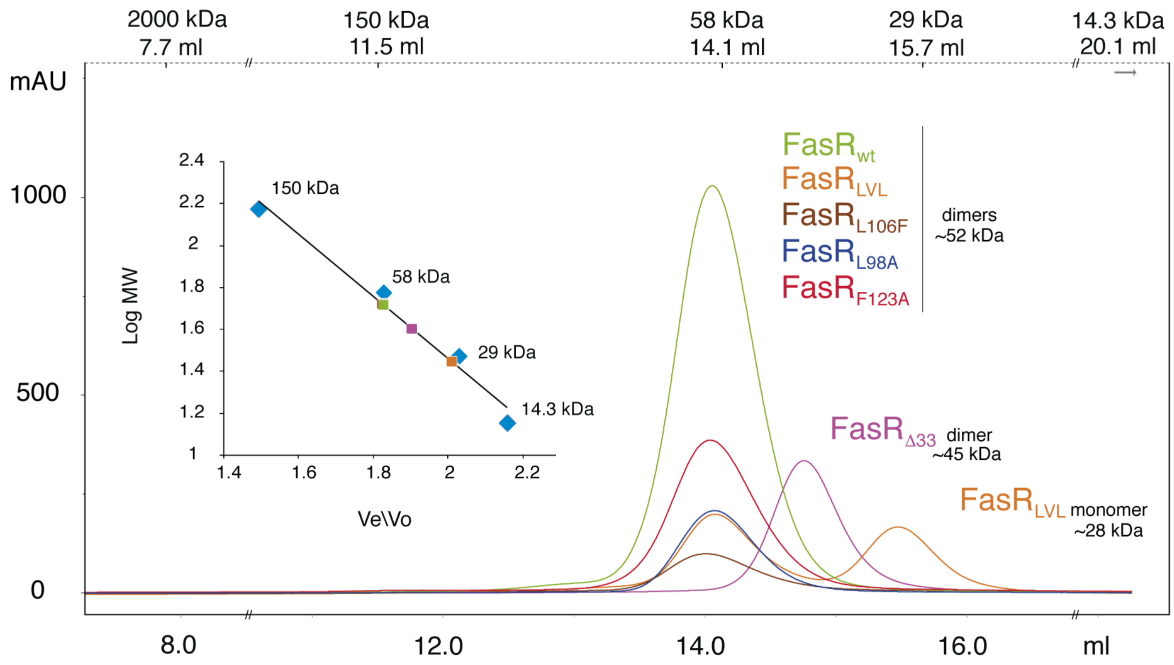


**Supplementary Figure S3**. Size exclusion chromatography purifications of FasR wild-type, Δ33 truncated and selected point-mutants (colour coded to match elution curves and inset calibration marks). Elution volume (Ve) is plotted on the x-axis, and 280 nm absorbance (in milliunits) on the y-axis. The inset shows a calibration curve with globular molecular weight (MW) standards (blue squares). The three different elution volumes are extrapolated to predict apparent MW of the different species. The top x-axis indicates the corresponding elution positions of the 5 standard MW marker proteins. Note that the mutant $FasR_{LVL}$ is the only one that displayed an effect on the quaternary structure, with ~50% fractions of dimeric and monomeric species (only the peak corresponding to the dimer was recovered for further functional analyses by EMSA); for the rest, all proteins eluted as dimeric forms.

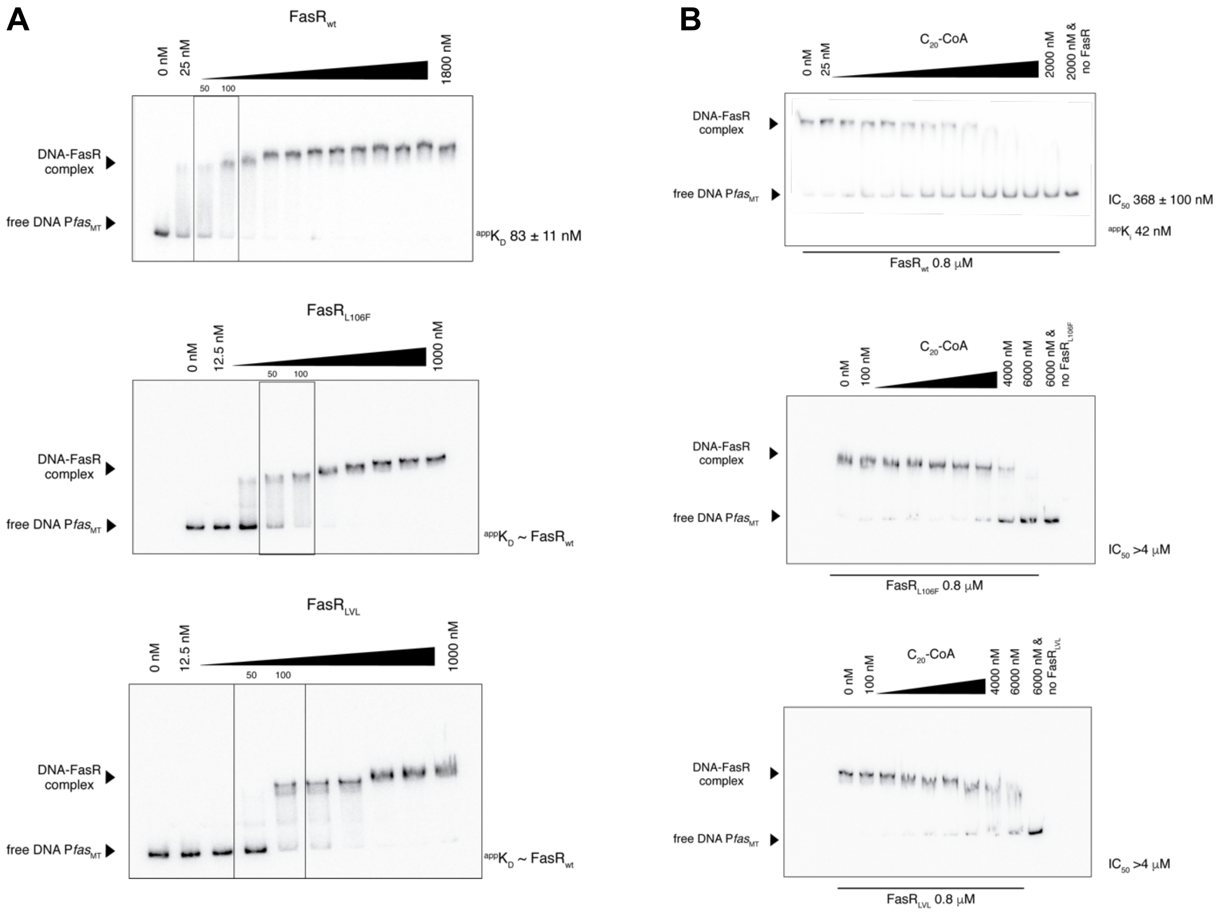


**Supplementary Figure S4.** Electrophoretic mobility shift assays comparing $FasR_{wt}$ vs the tunnel-occluding mutants $FasR_{L106F}$ and $FasR_{LVL}$. Representative gels are shown, all assays were done in duplicate or triplicate. **A.** increasing concentrations of FasR proteins were incubated with the P*fas*$_{MT}$ cognate DNA binding-site. Densitometry of band intensities allowed for quantification of $^{app}K_D$ affinity constants (see Methods for precise calculation procedures). The two mutants have similar DNA-binding behaviour than that of the wild-type, the boxes highlight the protein concentrations around which the $^{app}K_D$ constants fall in each case (values indicated on top). Triplicate assays for $FasR_{wt}$ allowed for standard error of the mean estimation, data for mutants were done in duplicate, their values are thus semi-quantitative. **B.** increasing concentrations of $C_{20}$-CoA effector were used to outcompete the different FasR variants (at fixed concentrations) complexed to cognate DNA. $IC_{50}$ and $^{app}K_i$ constants (see Methods for precise calculation procedures) were quantitated from the dose-response curves of effector concentration on DNA-binding affinity. Note the significantly reduced association of $C_{20}$-CoA to the mutants, barely allowing for any detectable effect. $^{app}K_i$s could not be reliably determined for the two FasR mutants, likely due to the very high $C_{20}$-CoA concentrations needed to start observing a dissociating effect (such concentrations of the ligand might result in detergent-like activity inducing erratic protein behaviour).

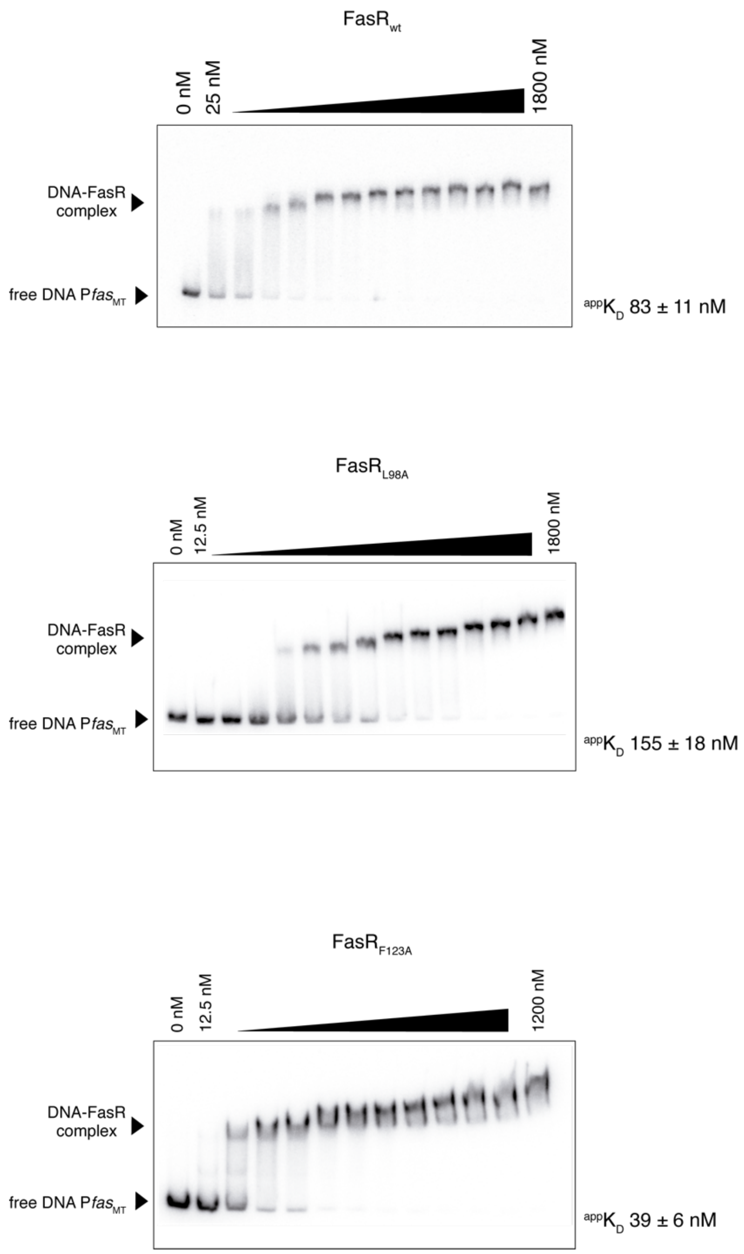


**Supplementary Figure S5.** Electrophoretic mobility shift assays comparing FasR$_{wt}$ vs the allosteric-uncoupling mutants FasR$_{L98A}$ and FasR$_{F123A}$. Densitometry of band intensities allowed for quantification of $^{app}K_D$ constants (see Methods for precise calculation procedure), a direct measure of protein:DNA affinity.

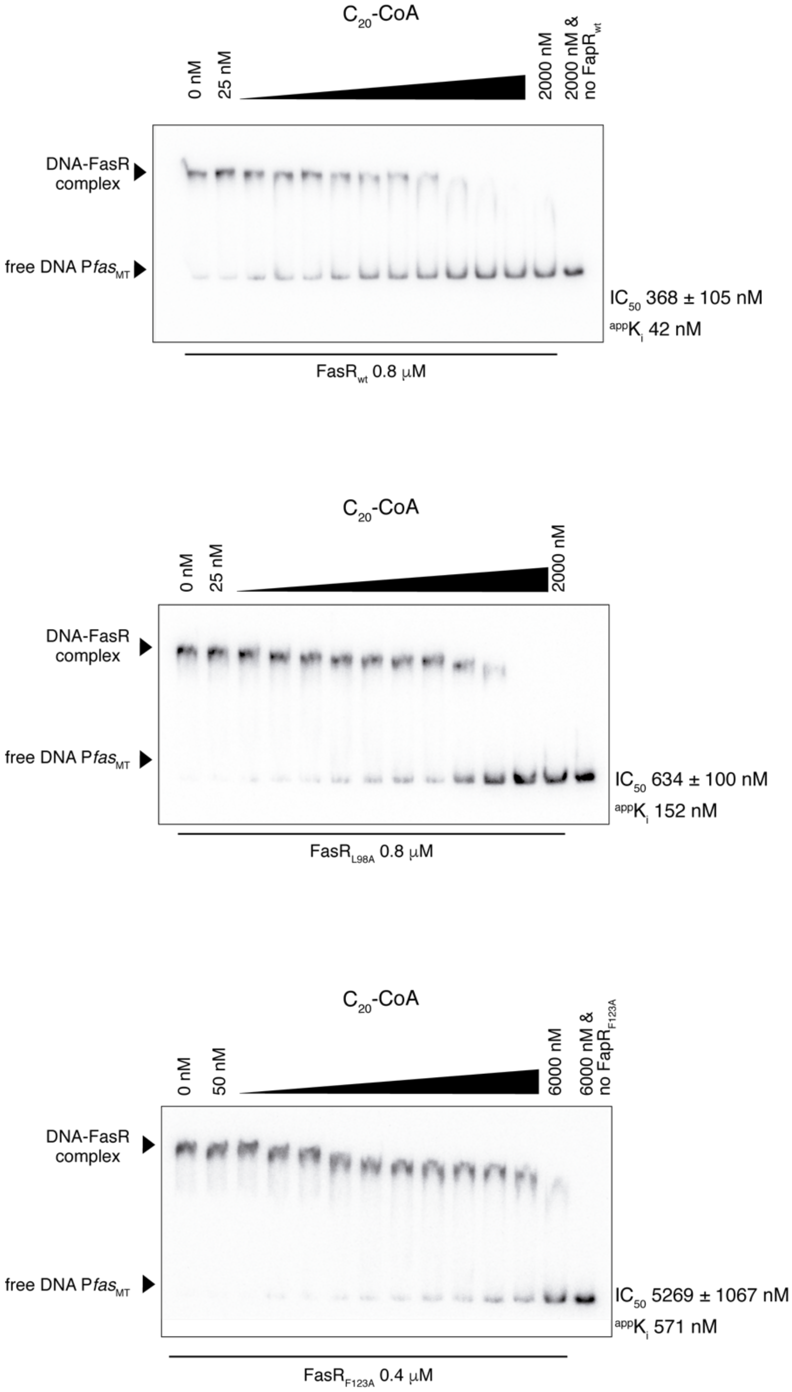


**Supplementary Figure S6.** Electrophoretic mobility shift assays comparing $FasR_{wt}$ vs the allosteric-uncoupling mutants $FasR_{L98A}$ and $FasR_{F123A}$. $IC_{50}$ and ${}^{app}K_i$ constants were quantitated from the dose-response curves of $C_{20}$-CoA effector concentration on DNA-binding affinities.

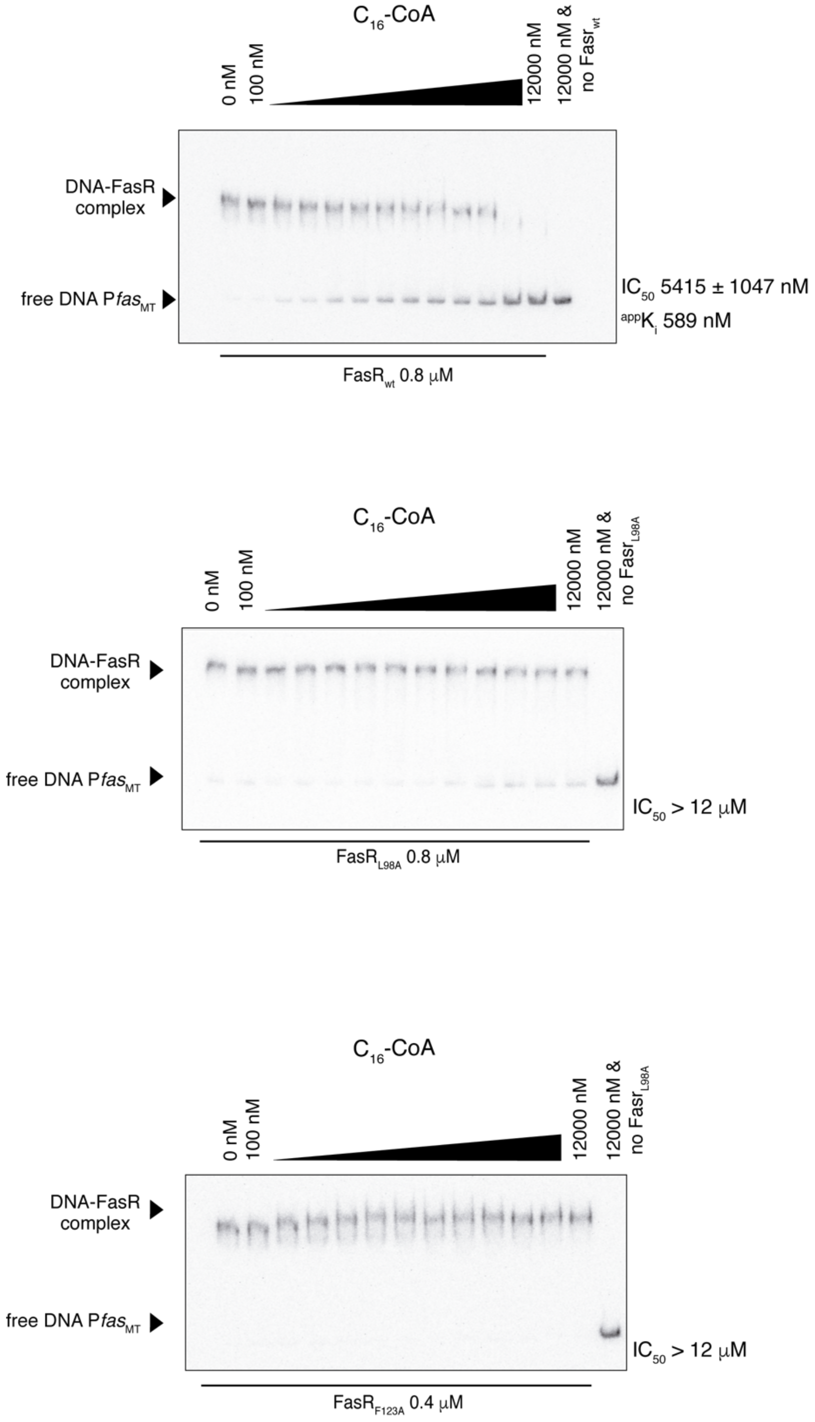


**Supplementary Figure S7.** Electrophoretic mobility shift assays comparing $FasR_{wt}$ vs the allosteric-uncoupling mutants $FasR_{L98A}$ and $FasR_{F123A}$. $IC_{50}$ and $^{app}K_i$ constants were quantitated from the dose-response curves of $C_{16}$-CoA effector concentration on DNA-binding affinities. Note that $^{app}K_i$s could not be reliably determined for the two FasR mutants, likely due to the very high $C_{20}$-CoA concentrations needed to start observing any effect.

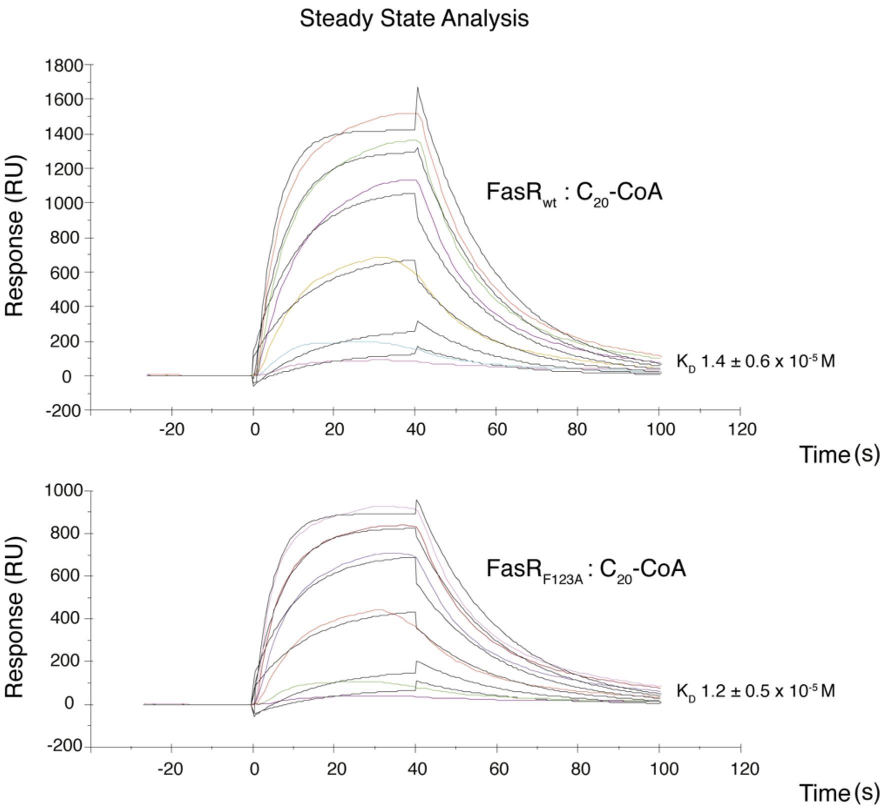


**Supplementary Figure S8**. Surface plasmon resonance assays. The binding of $C_{20}$-CoA to both FasR$_{wt}$ (top panel) and FasR$_{F123A}$ (bottom panel) are similar, plotted in sensorgrams with arbitrary SPR response units (y-axis) *vs* time (x-axis), with each curve representing increasing analyte concentrations ($C_{20}$-CoA). Dissociation constants ($K_D$) are calculated from the measured kinetic association and dissociation constants ($K_D = k_{off}/k_{on}$).

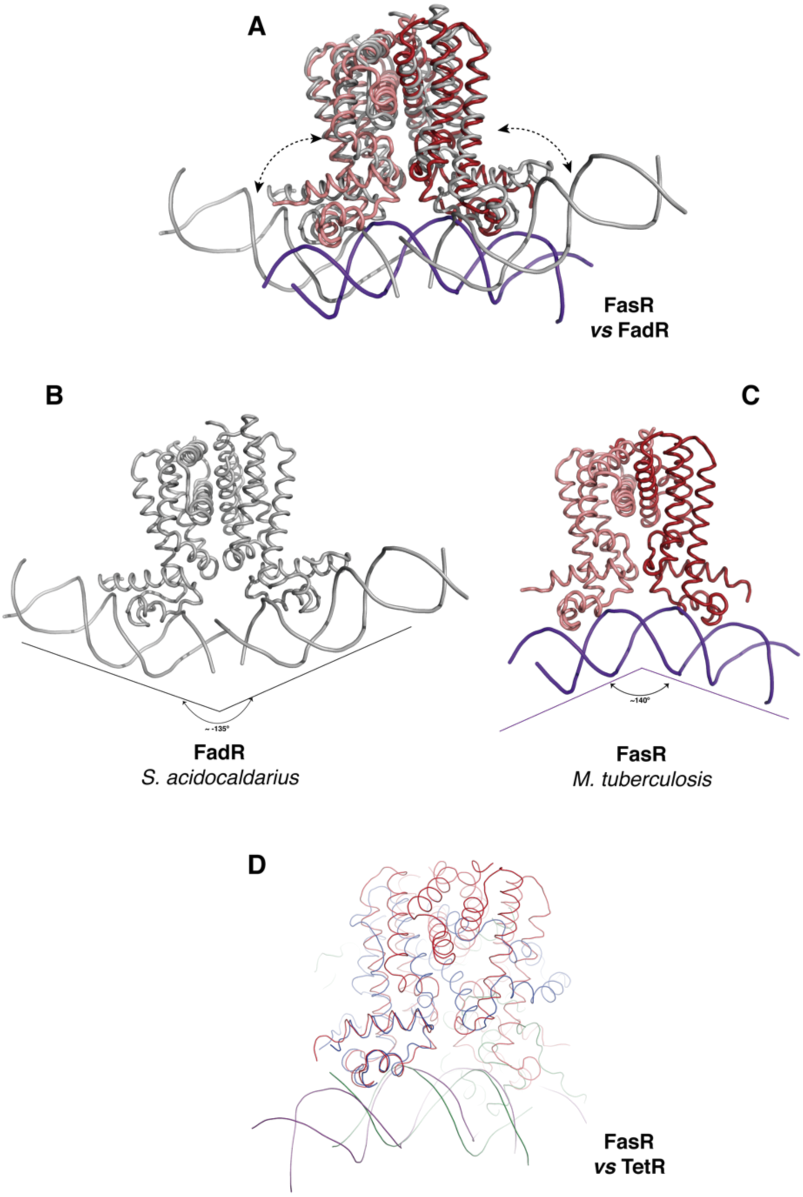


**Supplementary Figure S9. Structural comparison of *M. tuberculosis* FasR:DNA complex and *Sulfolobus acidocaldarius* FadR:DNA (pdb 6EN8). A.** Superposition of both structures, minimizing the rmsd between the effector binding domains of both dimers (4.3 Å rmsd for 250 C$\alpha$ atoms aligned). Only one of the FadR dimers in the asymmetric unit was used (6EN8 has three dimers in the ASU, the other two lie on straight segments of the DNA fragment). Note the significant shift on the positions of the HTH DNA-binding domains. **B.** Same perspective as in panel a, showing only the FadR:DNA complex. The kink angle on the DNA is indicated. **C.** Same perspective as in panel a, showing only the FasR:DNA complex. The kink angle on the DNA is indicated, and has an opposite value compared to FadR. **D.** FasR:DNA (in red:purple) is superposed onto TetR:DNA from *E. coli* (in blue:green), highlighting the similar bending angle on the DNA, despite important structural differences in the dimeric effector-binding core.

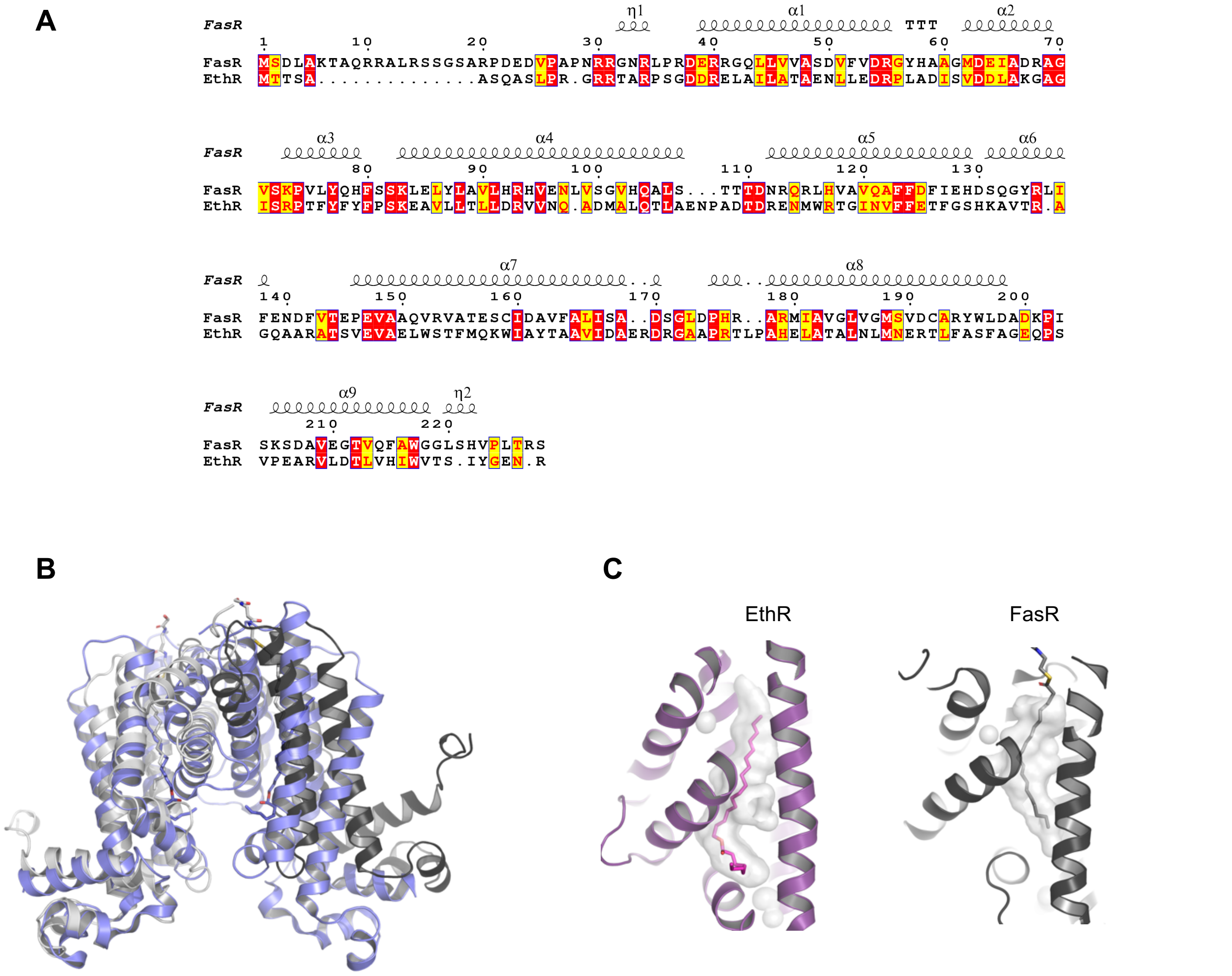


**Supplementary Figure S10. Detailed analysis comparing FasR and EthR from *M. tuberculosis*. A.** Pairwise sequence alignment, with the secondary structure elements of FasR depicted on top of the sequences. Identical residues are shown in white on red background, and physicochemicallly similar are shown in red on yellow background. **B.** Structural superposition by minimizing shifts of the main chain atoms of one protomer (to the left) of both regulators, FasR in tones of grey and EthR (pdb 1U9N) in slate blue. Ligands are depicted in sticks and coloured by atom, with carbons matching the corresponding cartoon tones. Note how the relative position of the second protomer results quite shifted. As well, the distance between both protomers delimiting an inter-protomer cavity toward the bottom of the figure, is much narrower in EthR. **C.** Side-by-side comparison of the cavities(3), rendered in semi-transparent grey surfaces, highlighting the effector-binding tunnels of EthR (left side; PDB 1U9N, protein cartoon in purple, and hexadecyl octanoate ligand in sticks coloured by atom) and FasR (right side; protein cartoon in dark grey, and $C_{20}$-CoA in sticks coloured by atom).

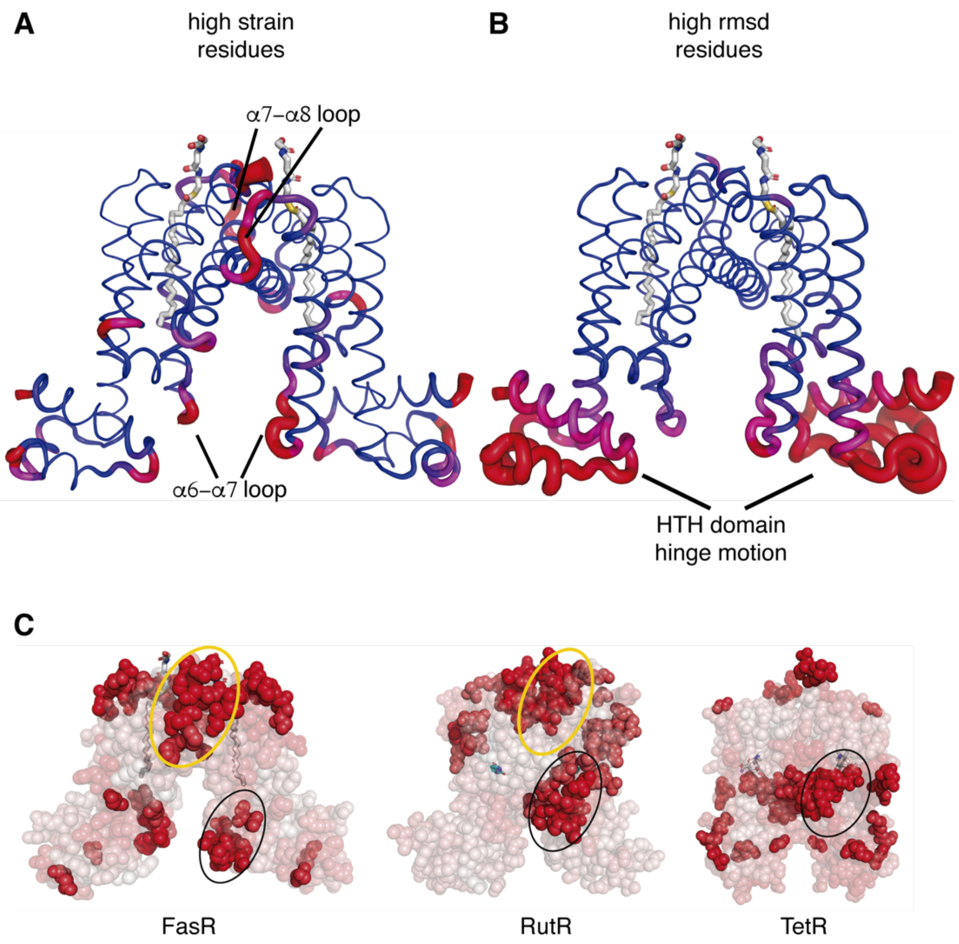


**Supplementary Figure S11. Strain analysis. A.** The measurement of mechanical strain per residue is mapped onto the ribbon representation of FasR$_{\Delta33}$-C$_{20}$-CoA with a blue-magenta-red colour ramp (from lower to higher strain values) and with the radius of the ribbon tube. Two loops with highest figures are indicated. Similar results were obtained comparing FasR$_{\Delta33}$-C$_{20}$-CoA vs FasR$_{\Delta33}$-C$_{14}$ or FasR$_{\Delta33}$-C$_{20}$-CoA vs FasR-DNA. **B.** The same representation and view perspective as in panel (a), now plotting the root mean squared deviations per residue. Once again, normalized according to the rmsd range in each case, a similar pattern is obtained comparing FasR$_{\Delta33}$-C$_{20}$-CoA vs FasR$_{\Delta33}$-C$_{14}$ or FasR$_{\Delta33}$-C$_{20}$-CoA vs FasR-DNA. **C.** The strain analysis mapping shown in panel (a) is shown here in semi-transparent sphere representation, with a white-to-red colour ramp (from lower to higher strain values). Residues with highest strain scores are depicted in solid spheres. Comparison with RutR and TetR, chosen here as two additional representative cases, highlights that loops $\alpha$6-$\alpha$7 (encircled with black lines) and $\alpha$7-$\alpha$8 (orange lines, only in FasR and RutR) play a conserved role in bearing with most shear strain. Note that in all cases the

$\alpha$6-$\alpha$7 loop is contacting both the effector molecule, as well as the protein elements connecting effector-binding and DNA-binding domains.

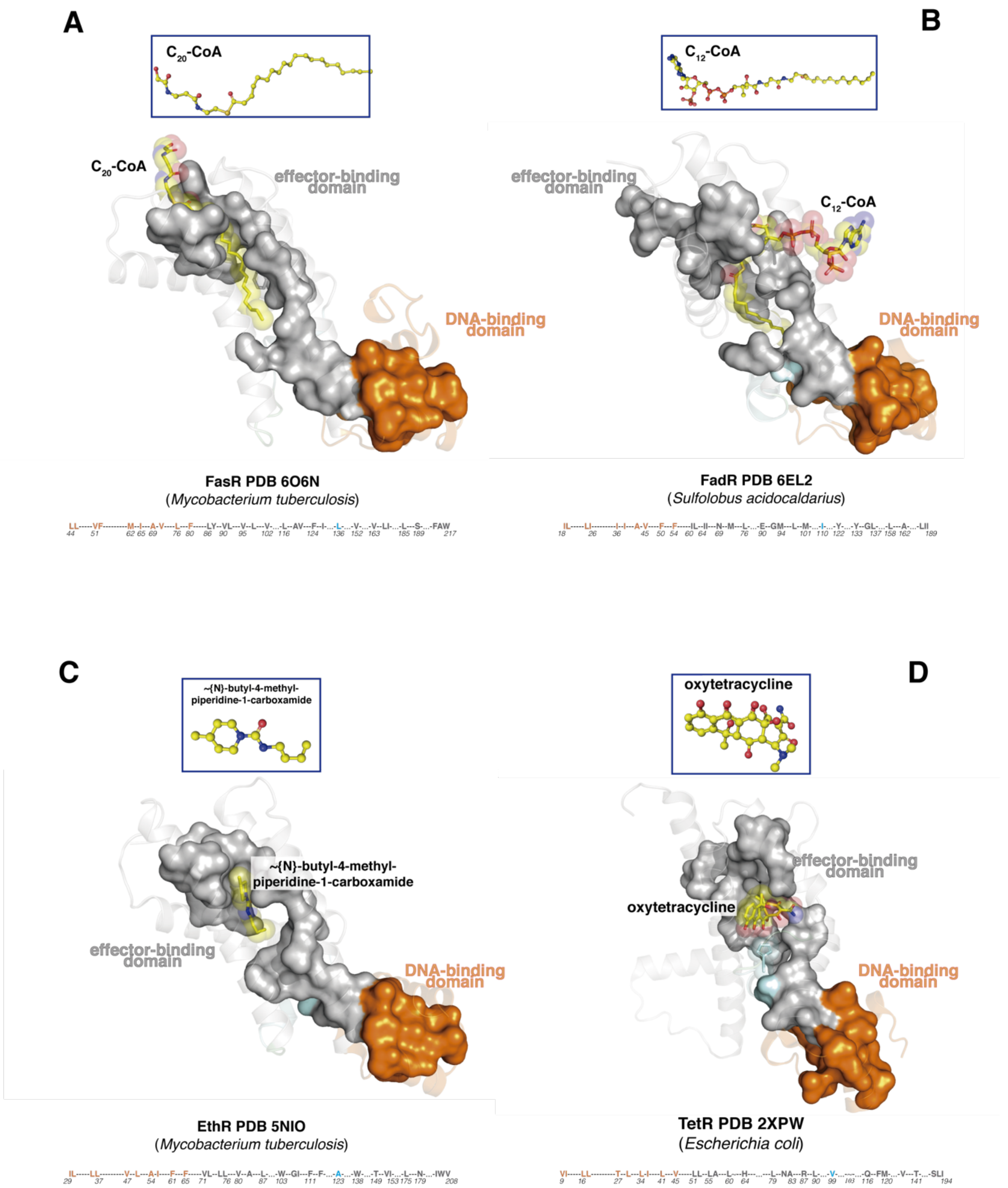


**Supplementary Figure S12. Hydrophobic spine conserved in TFRs.** The spine is continuous, connects both domains, and is completed by the binding of the effector ligand in all cases. **A-D.** Distinct TFRs (FasR, FadR, EthR, TetR) illustrate the conservation of the hydrophobic spine (spine residues are shown with their molecular surface), connecting effector- to DNA-binding domains. The effector-binding domain is coloured in grey, helix $\alpha6$ and the $\alpha6$-$\alpha7$ loop in cyan, and the DNA-binding domain in orange. The effector molecules in atom-coloured sticks are labelled, with transparent spheres overlaid (insets show their markedly disparate structures). Below each panel, the sequence of the spine is numbered according to each one of the  proteins' sequences (colours respect the domains' scheme).

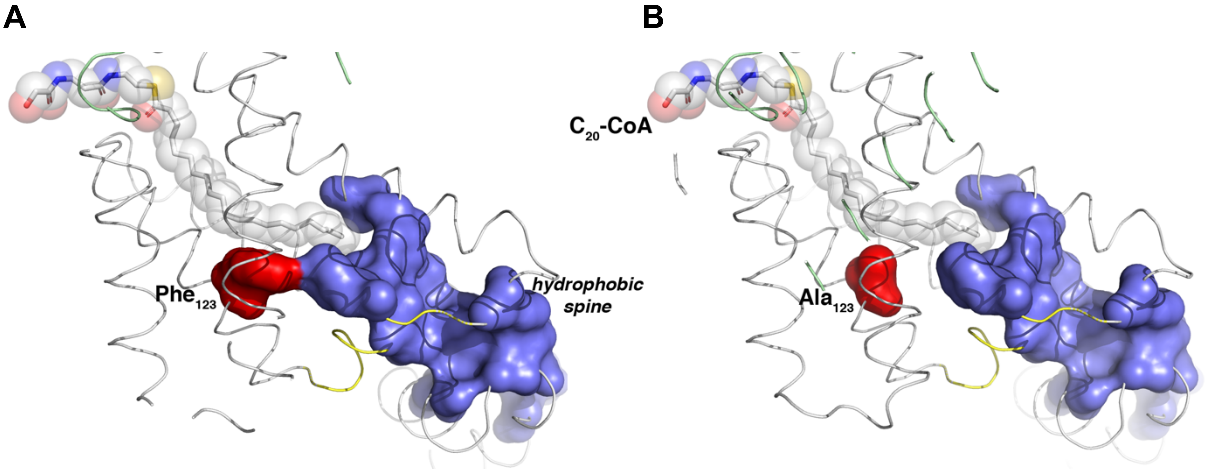


**Supplementary Figure S13. Uncoupling mutations in FasR likely break or destabilise the hydrophobic spine. A.** For clarity only the bottom portion of the hydrophobic spine is highlighted (blue molecular surface). The bound $C_{20}$-CoA effector molecule is shown as sticks, with overlaid transparent VDW spheres (coloured by atom type). Note the way by which the effector completes the hydrophobic spine. $Phe_{123}$ is highlighted in red surface. **B.** The substitution by $Ala_{123}$ destabilises the spine. Without the effector bound, the HTH domain could likely adopt an even floppier state, explaining higher DNA-binding affinity (see $^{app}K_D$ constants in Table 2). The effector can still bind with similar affinity (Supplementary Fig. S8), since the F123A substitution doesn't influence substantially. The hydrophobic spine not being stable enough, the HTH domains are still considerably flexible and DNA is able to accommodate inducing a closed conformation of the protein.

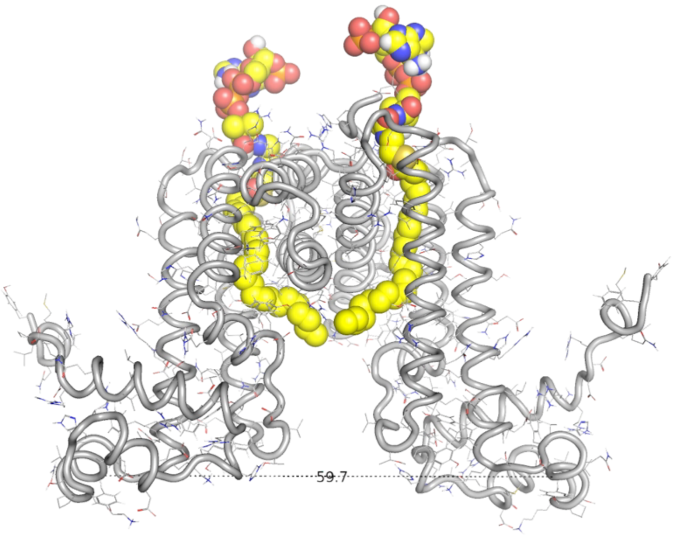


**Supplementary Video 1.** Molecular dynamics 10 ns trajectory. The animation was generated by sampling frames every 250 ps. The distance between the centres of mass of $Tyr_{77}$ on helix $\alpha3$ is indicated for all frames in this ensemble. The protein is represented as ribbons with amino acids as lines, and the $C_{26}$-CoA ligand is shown as spheres coloured by atom.

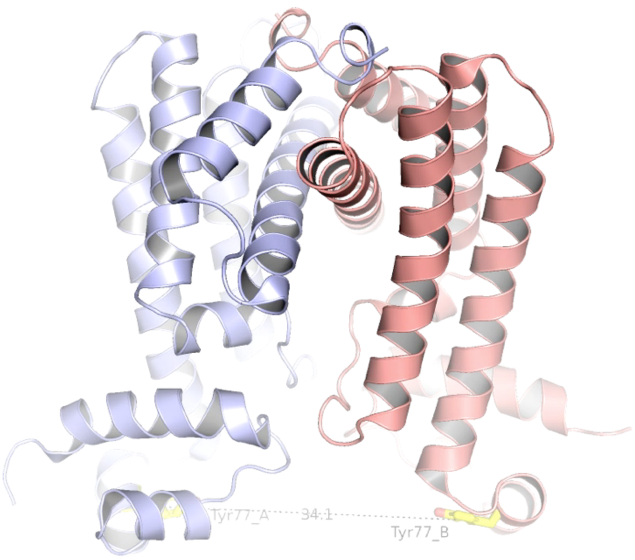


**Supplementary Video 2.** Linear morphing transforming the FasR-DNA crystal structure to the $FasR_{\Delta 33}$-$C_{20}$-CoA complex, looping back to finish on FasR-DNA. The distance between the centres of mas of $Tyr_{77}$ (shown as sticks coloured by atom) on helix $\alpha 3$ is indicated all along the animation. The protein is represented as cartoons with both protomers distinguished with colours. The $C_{20}$-CoA ligand as well as the DNA were not included to improve clarity.

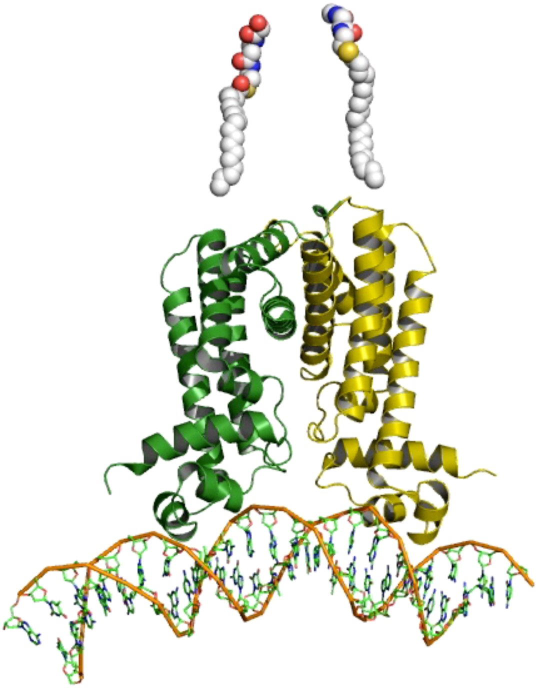

**Supplementary Video 3.** Linear morphing transforming the FasR-DNA crystal structure to the FasR$_{\Delta 33}$-C$_{20}$-CoA complex, looping back to finish on FasR-DNA. Important details of each structure are included: the action of acyl-CoA binding in stabilizing the HTH-open configuration; details of the tunnel; and association of FasR with the cognate DNA.

```
>UniRef100_H9ZSN6/1-191 Transcriptional regulator n=1 Tax=Thermus thermophilus JL-18 TaxID=798128
RepID=H9ZSN6_THETH
---------------------------------------------------------mg--TPTRTRILQE-
AAKLFTEKGYEATSVQDLAQALGLSKAALYHHF-G-S-KEEILYEIS-LLA-L----EGLVAA-GEKALE--
----------------------VADPK--E----AL-----RRF-ME--A----HARYFEE---NR----PF
---fVAM-L----QG------------L-QS--LS-P--EH--------REA----TV---RL-R----DR-
HEEN------L-------------------RA-IL---RRG-VE-Q--GVF-RE-VDVA--LAGRAVLSML-
N------W----------M-I-------RWF-R-P---------DGPMR----------------A-EEVAR
-AYHDLILRGLE-----rgsa--------------------------------------------------
--
>UniRef100_A0A191V4W6/1-215 TetR family transcriptional regulator n=1 Tax=Streptomyces parvulus TaxID=146923
RepID=A0A191V4W6_9ACTN
------------------------------------------------------marqlRAEQTRATIVDA-
AADLFDRHGYESTSLSEIVAHAGVTKGALYFHF-A-A-KEDLAHAIM-EIQ-S----RTLRRL-ADELTG--
-----------------R----GYTSL--E----AL-----MRI-TF--G----MARLCEE----G----PV
----LRA-G----LR------------L-AT--AG-V--PV--------RSP----LP---HP-F----TE-
WREI------A-------------------TS-RL---LDA-VR-Q--ADV-HPDIDVD--SVAHTLVCSV-
V------G----------T-Rl----vSGTL-E-P---------AGR-H----------------P-RR-LA
-QMWHILIRGMVP----vtrraryvtlaarleretasa----------------------------------
--
>UniRef100_D6ELN6/1-215 CprB n=1 Tax=Streptomyces lividans TK24 TaxID=457428 RepID=D6ELN6_STRLI
------------------------------------------------------marqlRAEQTRATIIGA-
AADLFDRRGYESTTLSEIVAHAGVTKGALYFHF-A-A-KEDLAHAIL-EIQ-S----RTS-RR-LAKDLD--
----------------gR----GYSSL--E----AL-----MRL-TF--G----MARLCVQ---GP-----V
----LRA-G----LR------------L-AT--AG-V--PV--------RPP----LP---HP-F----TE-
WREI------A-------------------TS-RL---LDA-VR-Q--SDV-HQDIDVD--SVAHTLVCSV-
V------G----------T-Rv----vGGTL-E-P---------AGR---------E--------P-R-RLA
-EMWYILIRGMVP----vtrraryvtlaarleqetgtt----------------------------------
--
>UniRef100_UPI0009A6AAAF/1-196 TetR family transcriptional regulator n=1 Tax=Halobacillus salinus TaxID=192814
RepID=UPI0009A6AAAF
--------------------------------------------------------mkk-NKPKYKQIIDA-
AVEVIAENGYHSSQVSKIAKKAGVADGTIYLYF-K-N-KEDILVSLF-QEK-M----GQFIEK-IEQETN--
-----------------S----RQTAE--E----KL-----LKL-VE--T----HFEQLSA---DH----HL
----AIV-T----QL------E-----L-RQ--SN-K--DL--------RQK----IN---SV-L----KP-
YLNV------I-------------------DA-II---SEG-VE-E--GLF-RPNLDRR--LVRQMIFGTL-
D------E----------T-V------TNWV-M-K---------EQRYN----------------I-VDQAH
-EVHSLIVHGLA-----rssq---------------------------------------------------
--
>UniRef100_UPI0001798FAF/1-218 transcriptional regulator of the TetR/AcrR family n=1 Tax=Thermobifida fusca (strain YX)
TaxID=269800 RepID=UPI0001798FAF
-------------------------------gmgqrsdsdhvmaeattdkrqghtrgrt-GNDRRTQIIKV-
ATELFREKGYYATSLDDIADRIGFTKPAIYYYF-K-S-KEDVLFAIV-NSI-V----DEALER-FHAIAA--
-----------------G----PGSPG--E----RI-----HAL-LV--E----HTRTILR---NL----DA
----NTL-F----YN------E----rG-L---LS-P--ER--------ERE----MR---KR-E----RE-
YTEI------M-------------------QR-LY---AEG-VA-T--GEL-L-DVDPT--VATATLLGAA-
I-----------------W-T------YRWY-D-P---------EGRLS----------------A-DEVVE
-QITRLLLNGYR-----rpa----------------------------------------------------
--
>UniRef100_E6SGQ0/1-200 Regulatory protein TetR n=1 Tax=Thermaerobacter marianensis (strain ATCC 700841 / DSM
12885 / JCM 10246 / 7p75a) TaxID=644966 RepID=E6SGQ0_THEM7
---------------------------------------------------------mag----REQEILDA-
ARKLFRQKGYYATTMQDIAEAVGLQKASLYHYI-R-S-KEALLLQVA-GET-M----RLFHAE-LDRIEA--
-----------------A----GGSYA--E----QL-----AAA-IR--A----HVRVVAE---HQ----ET
----LTV-L----FR------E-----S-HA--LP-P--EH--------AEK----VR---GE-T----RR-
```

**Supplementary Data 1. Multiple sequence alignment of 2591 sequences corresponding to different TetR-like transcription factors.** Only the first page is shown here. For the full alignment a separate text file is available as Supplementary Data 1, in Multiple Fasta format, for ready visualization by JalView or similar programs. All sequence names include the UniProt accession code in the form of UniRef100_*UNIPROT_CODE*/*sequence_included_in _alignement* followed by self-explanatory metadata about each protein. Note that the last six sequences correspond to selected TFRs with known 3D structure that we have used for in depth structural analyses (Fig. 6 and Supplementary Fig. S10).
